# In contrast to T_H_2-biased approaches, T_H_1 COVID-19 vaccines protect Syrian hamsters from severe disease in the absence of dexamethasone-treatable vaccine-associated enhanced respiratory pathology

**DOI:** 10.1101/2021.12.28.474359

**Authors:** Aileen Ebenig, Samada Muraleedharan, Julia Kazmierski, Daniel Todt, Arne Auste, Martina Anzaghe, André Gömer, Dylan Postmus, Patricia Gogesch, Marc Niles, Roland Plesker, Csaba Miskey, Michelle Gellhorn Serra, Angele Breithaupt, Cindy Hörner, Carina Kruip, Rosina Ehmann, Zoltan Ivics, Zoe Waibler, Stephanie Pfaender, Emanuel Wyler, Markus Landthaler, Alexandra Kupke, Geraldine Nouailles, Christine Goffinet, Richard J.P. Brown, Michael D. Mühlebach

## Abstract

Since December 2019, the novel human coronavirus SARS-CoV-2 has spread globally, causing millions of deaths. Unprecedented efforts have enabled development and authorization of a range of vaccines, which reduce transmission rates and confer protection against the associated disease COVID-19. These vaccines are conceptually diverse, including e.g. classical adjuvanted whole-inactivated virus, viral vectors, and mRNA vaccines.

We have analysed two prototypic model vaccines, the strongly T_H_1-biased measles vaccine-derived candidate MeV_vac2_-SARS2-S(H) and a T_H_2-biased Alum-adjuvanted, non-stabilized Spike (S) protein side-by-side, for their ability to protect Syrian hamsters upon challenge with a low-passage SARS-CoV-2 patient isolate. As expected, the MeV_vac2_-SARS2-S(H) vaccine protected the hamsters safely from severe disease. In contrast, the protein vaccine induced vaccine-associated enhanced respiratory disease (VAERD) with massive infiltration of eosinophils into the lungs. Global RNA-Seq analysis of hamster lungs revealed reduced viral RNA and less host dysregulation in MeV_vac2_-SARS2-S(H) vaccinated animals, while S protein vaccination triggered enhanced host gene dysregulation compared to unvaccinated control animals. Of note, mRNAs encoding the major eosinophil attractant CCL-11, the T_H_2 response-driving cytokine IL-19, as well as T_H_2-cytokines IL-4, IL-5, and IL-13 were exclusively up-regulated in the lungs of S protein vaccinated animals, consistent with previously described VAERD induced by RSV vaccine candidates. IL-4, IL-5, and IL-13 were also up-regulated in S-specific splenocytes after protein vaccination. Using scRNA-Seq, T cells and innate lymphoid cells were identified as the source of these cytokines, while *Ccl11* and *Il19* mRNAs were expressed in lung macrophages displaying an activated phenotype. Interestingly, the amount of viral reads in this macrophage population correlated with the abundance of Fc-receptor reads. These findings suggest that VAERD is triggered by induction of T_H_2-type helper cells secreting IL-4, IL-5, and IL-13, together with stimulation of macrophage subsets dependent on non-neutralizing antibodies. Via this mechanism, uncontrolled eosinophil recruitment to the infected tissue occurs, a hallmark of VAERD immunopathogenesis. These effects could effectively be treated using dexamethasone and were not observed in animals vaccinated with MeV_vac2_-SARS2-S(H).

Taken together, our data validate the potential of T_H_2-biased COVID-19 vaccines and identify the transcriptional mediators that underlie VAERD, but confirm safety of T_H_1-biased vaccine concepts such as vector-based or mRNA vaccines. Dexamethasone, which is already in use for treatment of severe COVID-19, may alleviate such VAERD, but in-depth scrutiny of any next-generation protein-based vaccine candidates is required, prior and after their regulatory approval.

## Introduction

In late 2019, a novel coronavirus (CoV) emerged causing severe acute respiratory disease (later termed COVID-19), that is regularly accompanied by cough, fever, and chest discomfort, and is fatal in a significant fraction of patients (Wu et al., 2020; Zhu et al., 2020). A novel beta CoV was identified as the causative pathogen, which was termed Severe Acute Respiratory Syndrome Coronavirus 2 (SARS-CoV-2). Since then, the virus caused a pandemic, resulting in nearly 274 million confirmed cases and 5.3 million deaths worldwide (as of Dec 21, 2021), with an overall case fatality rate of 2.03% (WHO, 2021). Remarkably, the first vaccine candidates were developed, tested in animal models, and approved for human use within the first year after pathogen identification. To date, more than 6.2 billion vaccine doses have been administered with 8 authorized vaccines based on three different vaccine technologies, namely mRNA vaccines, non-replicating adenoviral vector vaccines, or inactivated viruses.

Importantly, despite the remarkable pace of SARS-CoV-2 vaccine development, potential safety risks such as the induction of a more severe or altered clinical pathology after breakthrough infection in vaccinated patients were considered by most approaches. One possible risk associated with previous immunization in the context of respiratory infections is termed vaccine-associated respiratory disease (VAERD) (Munoz et al., 2021). For Dengue virus, induction of non-protective antibodies (Ab) by different serotypes or sub-potent vaccines has been correlated with sometimes dramatic disease enhancement by a process termed antibody-dependent enhancement (ADE) (Halstead and O’Rourke, 1977; Halstead, 1988; Katzelnick et al., 2017). Such enhancement of disease has also been well described in the context of formalin-inactivated respiratory syncytial virus (RSV) tested as vaccines for infants and young children in 1966 (FULGINITI et al., 1969; Kapikian et al., 1969; Kim et al., 1969). Vaccinated children that were seronegative before vaccination and were later exposed to pathogenic RSV developed an enhanced and atypical phenotype of clinical symptoms. The number of children requiring hospitalization was significantly increased for the vaccine cohort (FULGINITI et al., 1969; Kapikian et al., 1969; Kim et al., 1969), resulting in a small number of associated fatalities (Kim et al., 1969). A low-affinity and non-neutralizing Ab response provoked by the formalin-inactivated virus was found to be causative for enhanced disease (Polack et al., 2002). In addition, lung sections of the fatal cases that were examined post mortem displayed monocytic infiltration with massive excess of eosinophils into the affected lung tissue (Kim et al., 1969). Accordingly, T_H_2-biased T cell responses with eosinophil infiltration have been described in different animal models, in which the animals had been vaccinated with formalin-inactivated virus prior to RSV infection (Ruckwardt et al., 2019).

Moreover, inactivated CoV preparations show potential for VAERD in CoV-infected animals or animal models for highly pathogenic human CoV. In contrast to unvaccinated control kittens, cats immunized with a recombinant S protein-expressing vaccinia virus developed a more severe form of the disease and early death syndrome following infection with feline infectious peritonitis virus (FIPV), which was linked to a low amount of vaccine-induced neutralizing antibodies in the sera of affected animals (Vennema et al., 1990). This enhancement of clinical pathology can be reconstituted by passive transfer of antibodies directed against the same virus strain before infection, and was correlated to accelerated virus uptake by macrophages via Fc receptor (Olsen et al., 1992; TAKANO et al., 2008). Such processes are considered a marker and assumed to be relevant for classical ADE as described for dengue virus (DENV) (Beltramello et al., 2010; Alwis et al., 2014; Katzelnick et al., 2017). Additionally, for the first two highly pathogenic beta-CoVes identified in human patients, SARS-CoV and MERS-CoV, VAERD was observed in animal models. Specifically, entry receptor-transgenic K18-ACE2 or hDPP4-mice that were immunized with whole inactivated virus vaccines developed severe immunopathology in lung tissue with infiltration of eosinophils after infection with SARS-CoV or MERS-CoV, respectively (Bolles et al., 2011; Tseng et al., 2012; Iwata-Yoshikawa et al., 2014; Honda-Okubo et al., 2015; Agrawal et al., 2016).

While the predictive value of these animal models remains to be scrutinized, this scenario prompted the scientific community and vaccine developers to carefully evaluate the predisposition of COVID-19 infection for disease enhancement and to avoid induction of T_H_2-biased immunity or non-functional Ab responses by the different front-runner vaccine candidates (Anderson et al., 2020; Corbett et al., 2020b; Corbett et al., 2020a; Jackson et al., 2020; Polack et al., 2020; Ramasamy et al., 2020; Walsh et al., 2020b; Sadoff et al., 2021; Stephenson et al., 2021; van der Lubbe et al., 2021). Accordingly, no ERD or ADE has been reported in vaccinated individuals to date. However, a recent publication described evidence for VAERD in a mouse model using a mouse-adapted recombinant SARS-CoV-2 (MA10) after immunization with alum-adjuvanted, whole-inactivated SARS-CoV-2, or adjuvanted heat-inactivated S protein. Following virus challenge, these animals developed considerable pneumonia with infiltration of eosinophils and neutrophils, a clear sign of VAERD, which was absent in naïve mice or mice immunized with a T_H_1-biased mRNA vaccine (DiPiazza et al., 2021).

We had previously described a measles vaccine-derived prototypic COVID-19 vaccine candidate which protected against infection with the low passage SARS-CoV-2 human patient isolate MUC-1 in the Syrian hamster (*Mesocricetus auratus*) model. In contrast, an alum-adjuvanted, recombinantly expressed, non-stabilized Spike protein, as model for a T_H_2-biased vaccine concept, failed to protect upon immunization despite induction of S-specific binding antibodies (Hörner et al., 2020).

Here, we report reproducible VAERD in hamsters vaccinated with alum-adjuvanted S protein. VAERD became evident from histopathological analyses of infected hamster lungs. *In situ* hybridization, immunohistochemistry, staining for eosinophils, and general histopathology revealed exaggeration of pneumonia and eosinophilic infiltration, while virus load was reduced compared to MOCK-immunized, infected animals. These effects could be correlated to a broadly enhanced dysregulation of gene expression, but especially induction of T_H_2-biased immune cells after vaccination mediating pathologic responses via T_H_2-marker cytokines interleukin (IL)-4, IL-5, and IL-13 most likely supported by IL-19 signalling as well as up-regulation of the major eosinophil attractant eotaxin-1 / CCL-11. Furthermore, using scRNA-Seq we attributed this dysregulation to specific immune cell subsets suggesting that ADE via Fc receptor-mediated skewing of virion uptake drives cytokine secretion by macrophages. Eosinophil infiltration and pathology observed in Alum+S vaccinated animals were dramatically reduced by treatment of infected animals with dexamethasone, while the whole syndrome was absent in animals that received our prototypic T_H_1-biased MeV model vaccine candidate.

## Methods

### Cells

Vero (African green monkey kidney; ATCC# CCL-81) and Vero clone E6 (ATCC# CRL-1586) cells were purchased from ATCC (Manassas, VA, USA) and cultured in Dulbecco’s modified Eagle’s medium (DMEM, Sigma Aldrich, Steinheim, Germany) supplemented with 10% Fetal bovine serum (FBS; Sigma Aldrich) and 2 mM L-glutamine (L-Gln, Sigma Aldrich). Cell cultures were incubated at 37°C in a humidified atmosphere containing 6% CO_2_ up to 30 passages after thawing of the initial stocks.

### Viruses

MeV_vac2_-SARS2-S(H) (Hörner et al., 2020) and MV_vac2_-GFP(P) (i.e. MV_vac2_-ATU(P) (Del Valle et al., 2007) with GFP inserted in the ATU (Malczyk et al., 2015) have been described previously. Subsequent passages were generated after TCID_50_ titration of infectious virus according to the method of Kaerber and Spaerman (Kärber, 1931). Stocks were generated by infection of Vero cells at an MOI = 0.03, and viruses in P3 or P4 were used for vaccination experiments. SARS-CoV-2 isolate MUC-IMB1 (Böhmer et al., 2020) was used in passage 3 on Vero-E6 cells after isolation from the patient as described before (Hörner et al., 2020).

### Animal experiments

All animal experiments were carried out in compliance with the regulations of German animal protection laws and as authorized by the RP Darmstadt and reported according tio the ARRIVE guidelines. Six to 12-weeks old Syrian hamsters (Envigo RMS, Venray, Netherlands) were randomized for age- and sex-matched groups. Animals were vaccinated intraperitonally (i.p.) in a prime-boost schedule (Days 0 and 21) with 5×10^5^ TCID_50_ of recombinant MeV-derived vaccine virus in 200 μl volume or subcutaneously (s.c.) with 10 μg recombinant SARS-CoV-2 S protein (Sino Biological Europe, Eschborn, Germany) adjuvanted with 500 μg aluminum hydroxide (Allhydrogel adjuvant 2%, vac-alu-250, InvivoGen, San Diego, CA, USA) in 100 μl volume. Blood was drawn on day 0 and 21 or 31. Splenocytes of vaccinated animals were isolated 14 days after second immunization or hamsters were challenge by intranasal application of 4 x 10^3^ TCID_50_ SARS-CoV-2 (isolate MUC-IMB1) in passage 3 in 100 μl volume. Animals were euthanized 4 days after infection.

### Virus neutralization test (VNT)

Virus neutralization tests (VNT) were performed as described previously (Hörner et al., 2020). In short, serum samples were diluted in 2-fold series in DMEM. 50 PFU MV_vac2_-GFP(P) or 100 TCID_50_ SARS-CoV-2 were mixed with diluted serum samples and incubated at 37°C for 1 h. Subsequently, the virus-serum mixture was added to 1 x10^4^ Vero or Vero E6 cells seeded 3 h before in 96 well plates (Thermo Fisher Scientific, Ulm, Germany). Cells were incubated for 4 days at 37°C in a humidified atmosphere containing 6% CO_2_. The virus neutralizing titer was determined as the reciprocal of the highest serum dilution that completely abrogated infectivity.

### IFN-gamma ELISpot analysis

Hamster interferon gamma (IFN-γ) enzyme-linked immunosorbent spot (ELISpot) analysis was performed using the Hamster IFN-γ ELISpot^BASIC^ kit (MABTECH, Nacka Strand, Sweden) in combination with multiscreen immunoprecipitation (IP) ELISpot polyvinylidene difluoride (PVDF) 96-well plates (Merck Millipore, Darmstadt, Germany) according to the manufacturer’s instructions. 5×10^5^ isolated splenocytes were co-cultured with different stimuli in 200 ml RPMI containing 10% FBS, 2 mM L-Gln, 10 mM HEPES pH 7.4, 50 mM 2-mecaptoethanol and 1% Penicillin-streptomycin. To re-stimulate SARS-CoV-2 specific T cells, isolated splenocytes were cultured with 10 mg/ml recombinant SARS-CoV-2 (2019-nCoV) Spike Protein (S1+S2 ECD, His tag) (Sino biological Europe). Recombinant Ovalbumin [10 mg/ml] served as negative protein control. In parallel, splenocytes were stimulated with 10 mg/ml MeV bulk antigen (Virion Serion, Würzburg, Germany). General stimulation of T cells was achieved using 5 mg/ml concanavalin A (ConA, Sigma-Aldrich) or recombinant 20 μg/ml Flagellin A produced in house (Schülke et al., 2011). Untreated splenocytes served as negative control. After 36 h of stimulation, cells were removed and plates were incubated with biotinylated detection antibodies and Streptavidin-HRP conjugate following the manufactures introductions using a 1 in 100 dilution for the streptavidin-HRP conjugate with 3-Amino-9-ethyl-carbazole (AEC; Sigma-Aldrich) dissolved in N,N-dimethylformamide (Merck Millipore) as substrate. Spots were counted using an Eli.Scan ELISpot scanner (AE.L.VIS, Hamburg, Germany) and analysis software ELI.Analyse V5.0 (AE.L.VIS).

### Determination of infectious virus lung titers

The right apical lobe of the lungs of infected animals was snap-frozen in liquid nitrogen and homogenized in 1 ml ice-cold DMEM containing 2 mM L-Gln and 1% Penicillin/Streptomycin in Lysing Matrix M tubes (MP Bioscience, Hilton, UK) using the Precellys24 tissue homogenizer (bertin TECHNOLOGIES, Montigny-le-Bretonneux, France) for 2x 10 sec at 6,000 rpm. Samples were kept on ice all time. Subsequently, organ debris was removed by centrifugation (13 min, 6,800 rpm, 4°C). Vero E6 cells were inoculated with the supernatants in a 10-fold dilution series for 7 d at 37°C. SARS-CoV-2 organ titer was calculated by the TCID_50_ method of Kaerber and Spearman according to virus-induced CPE and adjusted to 1 g of tissue.

### RNA preparation

The right middle lobe of the lungs of infected animals was homogenized in 1 ml TRIzol Reagent (Ambion, Thermo Fisher Scientific) in Lysing Matrix M tubes (MP Bioscience, Hilton, UK) using the Precellys24 tissue homogenizer (bertin TECHNOLOGIES) for 2x 15 sec at 6,000 rpm. Samples were kept on ice all the time. Organ debris was removed by subsequent centrifugation (13 min, 6,800 rpm, 4°C). Clear supernatant was used for RNA purification with Direct-zol RNA MiniPrep kit (Zymo research, Freiburg (Breisgau), Germany) according to the manufactures introduction.

### Quantitative reverse-transcription PCR (qRT-PCR)

RNA samples were quantified by quantitative reverse transcription-PCR (qPT-PCR) using Superscript III one step RT-PCR system with Platinum Tag Polymerase (Invitrogen, Darmstadt, Germany). Primer and probe sequences for mRNA encoding the SARS-CoV-2 E gene (Corman et al., 2020), hamster RPL18 (Zivcec et al., 2011), IL-4, and IL-13 (Espitia et al., 2010) were used as described. Primers for detection of Eoatxin-1 (Stanelle-Bertram et al., 2020) and forward primer sequences for IL-5 (Mendlovic et al., 2015) were ordered as described. The reverse primer sequence for IL-5 was adapted according to RNA Seq results of hamster lungs as described in this manuscript. Probes for mRNA encoding Eotaxin and IL-5 were designed as shown in Suppl. Tab. 3. Reactions were run in 96-well plates (Bio-Rad Laboratories, Hercules, CA) using CFX96 qPCR cycler (Bio-Rad Laboratories) and 5 μl RNA in a total reaction volume of 25 μl in triplicates. An internal Hamster reference (linear range, 4.5×10^6^ to 4.5×10^2^ copies, (Hörner et al., 2020)) was used for quantification of SARS-CoV-2 E gene copy numbers. This reference was validated for copy numbers of RPL18 housekeeping gene by utilization of a PCR product DNA reference generated as described (Osterrieder et al., 2020), and was used for quantification in subsequent runs (linear range, 1.8×10^5^ to 1.82×10^2^ copies). Following cycling conditions were used for analysis of all analyts: reverse transcription for 10 min at 55°C, denaturation for 180 sec at 94°C, followed by 45 cycles of 15 sec at 94°C and 30 sec at 58°C. Quantified sample copy numbers were normalized to copy numbers of the hamster housekeeping gene RPL18. If direct quantification of the analyts was not possible, the ΔΔC_t_ method was used.

### Total RNA Seq

The isolated RNA samples were used for NNSR priming based RNA-Seq library preparation (Levin et al., 2010)) as described in (Brown et al., 2020), vRNA NGS section) with the following modifications. Total RNA, was used for rRNA removal using the QIAseq FastSelect –rRNA HMR Kit (Qiagen) in combination with reverse transcription as follows. A 35 μl reaction mixture containing 1 μg RNA, 100 pmol NNSR_RT primer (gctcttccgatctctNNNNNN), 8 μl of 5x SuperScript IV buffer (Invitrogen) 20 pmol dNTPs and 1 μl of FastSelect-rRNA mix was subjected to the following hybridization protocol: 75°C 2 min, 70°C 2 min, 65°C 2 min, 60°C 2 min, 55°C 2 min, 37°C 5 min, 25°C 5 min, store at 4°C. For cDNA synthesis the reaction above was supplemented with dithiothreitol (10 mM), 20 units of RiboLock ribonuclease inhibitor (Thermo Fisher Scientific) and 200Us of SuperScript IV reverse transcriptase in a final reaction volume of 40 μl and incubated 45°C 5 min, 70°C 15 min.

The smears of 200-500 base pairs of the final barcoded libraries were purified from a 1.5% agarose gel and sequenced on a NextSeq 550 Illumina instrument using a single-end 86 bp setting. The RNS-Seq library preparation method used results in reads that start with the same two initial nucleotides. Hence, these were removed when performing quality- and minimum length-read trimming with the fastp algorithm (Chen et al., 2018)with the default parameters.

### Isolation of single cells from lung tissue

The right caudal lope of the lung was removed from the body and transferred on ice in PBS containing 1% BSA (w/v) and 2 mg/ml Actinomycin D (Sigma-Aldrich) for further processing. Lung tissue was incubated in 2 mL Dispase (Corning, Bedford, MA, USA) containing 2 mg DNAse (AppliChem, Darmstadt, Germany), 4.6 mg Collagenase B (Roche, Basel, Switzerland) and 2 mg/ml Actinomycin D (Sigma-Aldrich) and at 37 °C for 30 min. Digest was stopped by the addition of cold PBS containing 0.5% (w/v) BSA and Actinomycin D. The tissue was disrupted by pipetting in a repeated pumping motion. Cell suspension was collected and filtered through 70 mm-filter to obtain a single-cell suspension. Red Blood cell were lysed by incubation with RBC lysis buffer (Santa Cruz Biotechnology, Dallas, Texas, USA) for 4 min at room-temperature. Lysis reaction as stopped by the addition of PBS containing 0.04% BSA, the supernatant was removed by centrifugation (1,200 rpm, 6 min, 4 °C) and cells were resuspended in PBS containing 0.04% BSA. Barcoding of single cells and RNA Isolation was performed using the Chromium controller and Chromium Next Gem Chip G Single Cell Kit and Chromium Next Gem Single cell 3’ GEM, Library & Gel Bead Kit v3.1 (10x Genomics B.V., Leiden, The Netherlands) according to the manufactures instructions.

### Single-cell RNA-Seq

After enzymatic fragmentation and size selection, resulting double-stranded cDNA amplicons optimized for library construction were subjected to adaptor ligation and sample index PCRs needed for Illumina bridge amplification and sequencing according to the manufacturer’s instruction (10x genomics). Single cell libraries were quantified using Qubit (Thermo Fisher) and quality-controlled using the Bioanalyzer System (Agilent). Sequencing was performed on a Novaseq 6000 (Illumina), aiming for 200 Million reads per library (read1: 28, read2: 150 nucleotides). Data were analysed using CellRanger v5.0 (10X Genomics) using hamster and SARS-CoV-2 genome scaffolds, and the R packages Seurat v4.0 (Hao et al., 2021) and DoRothEA v3.12 (Holland et al., 2020) were used for cell clustering, annotation, and transcription factor activity analysis. Median gene number detected per cell ranged between 2000 and 4400, with 3800–18500 median UMI counts per cell. Gene set variation analysis (GSVA) was performed using the GSVA R package (Hänzelmann et al., 2013) and gene set enrichment analysis was performed using the clusterProfiler R package (Wu et al., 2021).

### Histopathology

The left lung lobe was carefully removed and immersion-fixed in 10% neutral-buffered formalin for 7 days. The tissue was subsequently paraffin-embedded and sections of 4 μm were prepared.

### Hematoxilin-Eosin staining

Hematoxylin–eosin staining was carried out in accordance with standard procedures (Mulisch and Welsch, 2015). H&E stained slices were subjected to histopathologic analyses on blinded samples.

### Sirius Red staining

For Sirius Red staining of lung tissue sections we followed the protocol published by Llewellyn with some modifications (Llewellyn, 1970). Shortly, sections were placed in Papanicolaous solution 1b Hematoxylin S (Sigma Aldrich) for 2 min and rinsed afterwards with water followed by ethanol, 3% HCl in ethanol and 70% ethanol. Subsequently, sections were stained for 90 min in alkaline Sirius red (0.5 g Direktrot 80, Sigma in 50% ethanol containing 0.1 ‰ NaOH) before rinsing with water. Sections were dehydrated afterwards with increasing ethanol concentrations and xylene. Finally, sections were covered with Entellan (Merck KGaA, Darmstadt, Germany).

### *In situ* hybridisation

To detect viral RNA in the lungs, fixed and routinely paraffin-embedded tissue sections were mounted on glass slides and analyzed by *in-situ* hybridization as described before (Halwe et al., 2021; Tscherne et al., 2021). For this, the RNAscope® 2.5 HD Assay – RED Kit (Bio-Techne, cat. no. 322360) was used according to the manufacturer’s instructions. Slides were incubated at 60 °C, deparaffinized with xylene and 100 % ethanol and pretreated with RNAscope® Pretreatment Reagents (cat. no. 322330 and 322000), to enable access to the target RNA. Subsequently, the RNA-specific probe, targeting the S protein of the SARS-CoV-2 virus (cat. no. 848561) was hybridized to the RNA. After the amplification steps, Fast Red substrate was administered to the samples for signal detection. Slides were counterstained with Gill’s Hematoxylin I and 0.02 % ammonia water. A RNAscope® Negative Control Probe (cat. no. 310043) was used in parallel to control background staining.

### Immunohistochemistry

For SARS-CoV-2 antigen detection, a monoclonal Ab against the nucleocapsid protein (clone 4F3C4, Ref: (Bussmann et al., 2006)) was used according to standardized procedures of avidin-biotin-peroxidase complex-method (ABC, Vectastain Elite ABC Kit, Burlingame, CA, USA). Briefly, 2-3 μm sections were mounted on adhesive glass slides, dewaxed in xylene, followed by rehydration in descending graded alcohols. Endogenous peroxidase was quenched with 3% hydrogen peroxide in distilled water for 10 minutes at room temperature. Antigen heat retrieval was performed in 10mM citrate buffer (pH 6) for 20 minutes in a pressure cooker. Nonspecific Ab binding was blocked for 30 minutes at room temperature with goat normal serum, diluted in PBS (1:2). The primary Ab was applied overnight at 4°C (1:50, diluted in TRIS buffer), the secondary biotinylated goat anti-mouse Ab was applied for 30 minutes at room temperature (Vector Laboratories, Burlingame, CA, USA, 1:200). Color was developed by incubating the slides with freshly prepared avidin-biotin-peroxidase complex (ABC) solution (Vectastain Elite ABC Kit; Vector Laboratories), followed by exposure to 3-amino-9-ethylcarbazole substrate (AEC, Dako, Carpinteria, CA, USA). The sections were counterstained with Mayer’s haematoxylin and coverslipped. As negative control, consecutive sections were labelled with an irrelevant Ab (M protein of Influenza A virus, ATCC clone HB-64). A positive control slide was included in each run. Slides were scanned using a Hamamatsu S60 scanner (Hamamatsu Photonics, K.K. Japan).

## Results

### Histological analysis and virus load in the lung

To study the efficacy of MeV-derived COVID-19 vaccines, we had vaccinated Syrian hamsters with the experimental vaccine candidate MV_vac2_-SARS2-S(H). Furthermore, we included a classic protein vaccine formulation consisting of recombinant, non-stabilized SARS-CoV-2 S protein, which revealed distinct changes in the composition of secondary structural elements (31.7% α-helical, 35.0% β-sheets vs. 19.3% α-helices and 50.8% β-sheets) compared to a stabilized soluble S by CD spectroscopy (Suppl. Fig. S1). Recombinant S was adjuvanted with aluminum hydroxide (Alum+S). The vaccinated animals were subsequently challenged in direct comparison with naïve or measles-vaccinated vector control hamsters (MV_vac2_-ATU(P)) using the low-passage SARS-CoV-2 patient isolate MUC-1 (as published in (Hörner et al., 2020) deposited as strain BavPat1. During the histological analysis of paraffin-fixed lung samples of these animals, the expected pathology was observed for the control animals in haematoxylin-eosin stained tissue slices of the left lobe of the lungs (Fig. 1A). In both naïve and MeV-control vaccinated hamsters, epithelia and endothelia in bronchii and vasculature, respectively, showed signs of inflammation, accompanied by hemorrhaging to different extents. Between 10 - 40% of the lung area had become dense due to cellular infiltrates, which consisted mainly of macrophages and lymphocytes; in general, no or few eosinophils or granulocytes were found in these samples. Single foci to moderate karyorrhexis was apparent in these areas (Suppl. Tab. 1). In samples of MeV_vac2_-SARS2-S(H)-vaccinated hamsters, the pathology of SARS-CoV-2 infection was significantly ameliorated, with reduced inflammation of epithelia, very few hemorrhages, only few eosinophils in perivascular regions, and no to minimal karyorrhexis. In contrast, animals vaccinated with alum-adjuvanted S protein displayed a considerably intensified pathogenesis: While bronchial epithelia showed few signs of inflammation, vascular endothelium revealed explicit inflammation. The area of dense infiltrates exceeded 50% of the investigated slice, with massive infiltration of eosinophils evident (Fig. 1A, C). Together, this histopathology suggests induction of vaccine-associated enhanced respiratory disease by the immunization of hamsters with Alum+S.

**Fig. 1:**
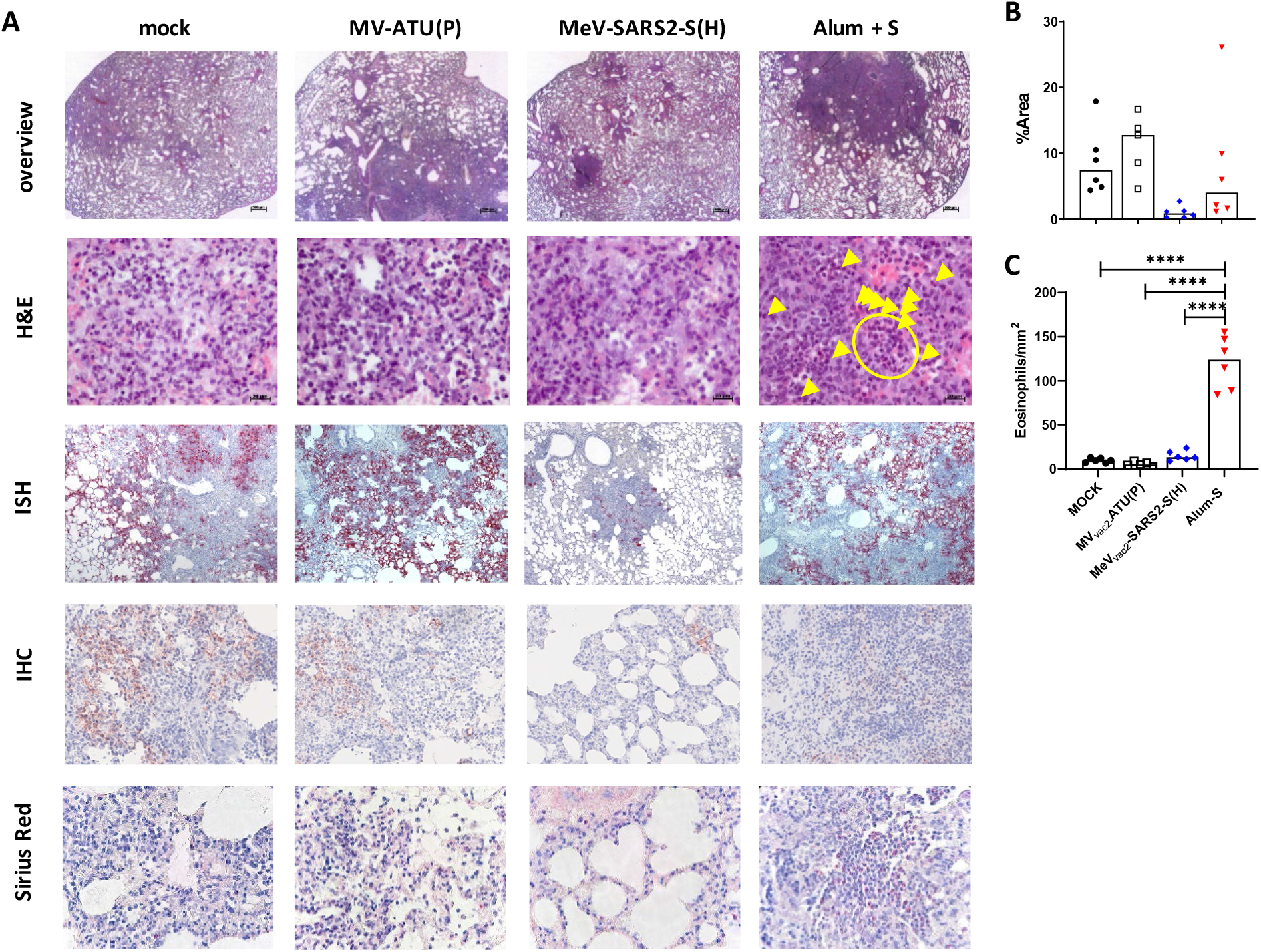
Pulmonary pathology in vaccinated Syrian hamsters upon challenge with SARS-CoV-2. **(A)** Depicted are representative sections of left lobes of vaccinated hamster lungs prepared 4 dpi with low-passage SARS-CoV-2. H&E staining (two top rows) reveals histopathological changes and immune cell infiltration. *In situ* hybridisation for SARS-CoV-2 RNA (ISH, 3^rd^ row) and immunohistochemistry staining for SARS-CoV-2 nucleocapsid protein (IHC, 4^th^ row) depict extent of infection, while sirius red staining (bottom row) reveals infiltration of eosinophils. Top row, 12.5x; other rows 400x magnification. **(B)** Quantification of infected tissue by determining the fraction of the slice are stained positive in ISH **(C)** Eosinophil infiltration was quantified for all animals is depicted. Each data point reveals the mean number of eosinophils per mm^2^ of individual animals. Yellow arrowheads and yellow circle depict individual eosinophils or clusters of eosinophils in H&E-stained samples, respectively.

The pathology correlated with SARS-CoV-2 infection as evident by *in situ* hybridisation (ISH) of SARS-CoV-2 RNA genomes (Fig. 1A third row, B) and immunohistochemistry for SARS-CoV-2 nucleocapsid protein N (Fig. 1A, fourth row). As expected from previous data, MV_vac2_-SARS2-S(H) vaccinated animals revealed significantly reduced staining for SARS-CoV-2 in both ISH and immunohistochemistry, and much less affected lung tissue, in general. These data are consistent with the previously published protective efficacy of this T_H_1-biased vaccine concept. In contrast, lungs of Alum+S vaccinated animals did not show amelioration of pathology. Despite somewhat reduced staining for SARS-CoV-2 genomes and N, inflamed areas in the lungs were larger than in naive animals. More impressive, H&E staining revealed in all samples massive infiltration of eosinophils into these inflamed areas. Sirius Red staining for eosinophils (Fig. 1A, bottom row) validated this finding, which became evident to this extent only in this vaccine cohort. Only in 3 out of 6 animals vaccinated with MeV_vac2_-SARS2-S(H), some eosinophils were found in perivascular regions of inflamed tissue (Suppl. Fig. S2). To quantify the extent of eosinophil infiltration and infection, we assessed the amount of infiltrating eosinophils in the different samples and the fraction of the slices’ area staining positive in ISH. Between 84 and 155 eosinophils/mm^2^ were found in lung sections of hamsters immunized with the alum-adjuvanted S, whereas in hamsters inoculated with MeV_vac2_-SARS2-S(H), or the MeV vector control, eosinophil numbers remained within the same range as uninfected control hamsters (3-24 eosinophils/mm^2^) (Fig. 1C). While 5 to 20% of the lung tissue stained positive for SARS-CoV-2 genomes in unvaccinated or measles-only vaccinated animals, lung tissue of hamsters vaccinated with MeV_vac2_-SARS2-S(H) was much less positive with a maximum of 5% of the area, while Alum+S vaccinated animals showed a split behaviour (Fig. 1B)

Thus, the COVID-19 hamster model established in our laboratory revealed features suggestive of vaccine-associated enhanced respiratory disease for animals vaccinated using the traditional T_H_2-biased (Hörner et al., 2020) alum-adjuvanted protein vaccine approach, but absence of such an effect for the T_H_1-biased measles-based COVID-19 vaccine candidate.

### Transcriptional profiling of SARS-CoV-2 infected hamster lungs with and without prior vaccination

To identify determinants underlying the differential degree of protection or pathology arising from T_H_1- or T_H_2-biased vaccination approaches, we performed RNA-Seq profiling of hamster lung tissue. Lung transcriptomes from SARS-CoV-2 infected hamsters were compared to infected animals previously immunized with either T_H_1- or T_H_2-biased vaccines. Numbers of differentially expressed genes (DEGs) in the lung were determined by comparison to baseline expression levels, derived from uninfected, unvaccinated control animals. Principal component analysis (PCA) of individual transcriptomes revealed segregation of signals according to infection and vaccination status, confirming distinct lung transcriptional responses to infection in the different groups (four animals per group, Fig. 2A). In infected lung tissue, significant transcriptional dysregulation was evident in the absence of vaccine-induced protection, with ~2,000 genes down-regulated and ~1,500 genes up-regulated in vaccine-naïve SARS-CoV-2 infected hamsters (Fig. 2B, left panel). However, prior vaccination with MeV_vac2_-SARS2-S(H) limited infection mediated changes in the lung transcriptional landscape, with a 40% reduction in numbers of significantly dysregulated genes. In line with this, lung resident viral RNAs were also 20-fold reduced when compared to unvaccinated animals (Fig. 2B, left panel) in accordance with the reduction of lung viral titers described earlier (Hörner et al., 2020). In contrast, challenge with SARS-CoV-2 after vaccination with alum-adjuvanted S protein resulted in comparable numbers of dysregulated genes to those observed in unvaccinated animals (Fig. 2B, left panel), with an increased median fold-change in gene expression also apparent (Fig. 2B, right panel). Vaccine-induced protection was also limited, with only a 2-fold reduction in lung-resident viral RNAs observed (Fig. 2B, left panel). Together, these data suggest prior immunization with a T_H_1-biased vaccine confers protection from disease, substantially reducing viral loads in hamster lungs, with a concomitant reduction in the magnitude of lung gene dysregulation. In contrast, the magnitude of lung gene dysregulation and associated viral load observed in hamsters immunized with a T_H_2-biased vaccine more closely resembled patterns seen in unvaccinated animals. However, inspection of the PCA plot revealed separate clustering of SARS-CoV-2 and Alum+S groups (Fig. 2A), indicating unique transcriptional signatures underlying Alum+S associated pathology.

**Figure 2:**
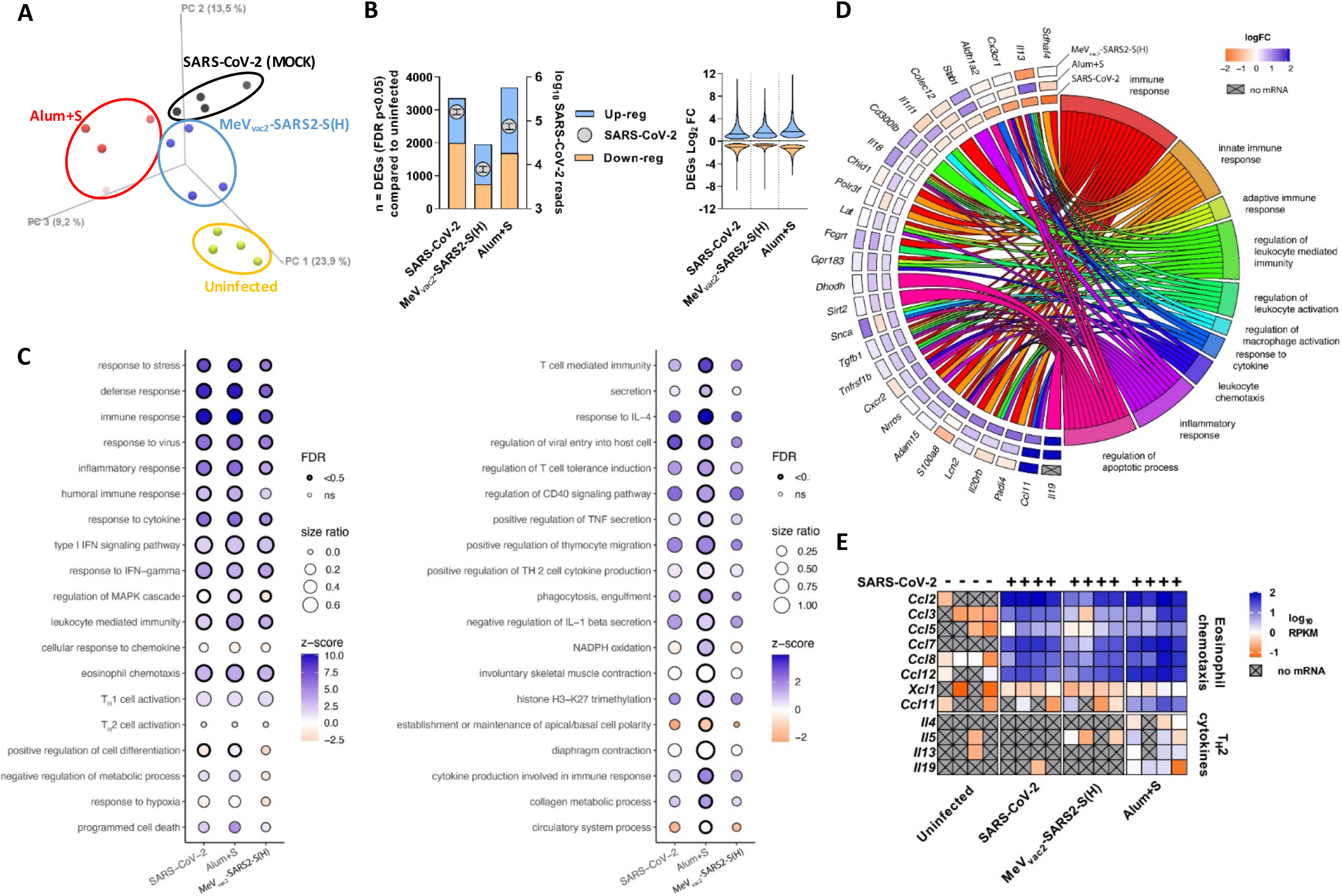
Global host gene dysregulation revealed by global RNA-Seq in lungs of infected, differently vaccinated hamsters. (**A**) Principle component analysis (PCA) of hamster lung transcriptomes. Transcriptomes from individual animals are represented by a single data point. Yellow, unvaccinated and not infected; blue, unvaccinated and infected; red, vaccinated with MeV_vac2_-SARS2-S(H) and infected; green, vaccinated with Alum-adjuvanted protein and infected). (**B**) Viral RNA copies and host gene dysregulation in hamster lungs after infection and vaccination. Left panel. Data plotted on the left y-axis represents the number of differentially expressed genes (DEGs) (FDR p-value <0.05) in the hamster lung under each condition, when compared to uninfected unvaccinated controls (n = 4). Blue: upregulated; orange: downregulated. Data plotted on the right y-axis (grey circles) represents the number lung-resident SARS-CoV-2 mapped reads per condition (mean±SEM). Right panel. Violin plots depict range of log_2_ fold-change in expression for all significantly dysregulated genes under each condition, when compared to uninfected unvaccinated controls. Horizontal lines, median values. (**C**) Gene ontology (GO) enrichment analysis of SARS-CoV-2 induced DEGs. GO categories are labelled on the y-axes. Circle size represents the ratio of significantly dysregulated genes relative to the total gene number in a specific GO term. Circles are shaded relative to activation z-score, with significantly enriched categories highlighted. Left panel. Selected GO categories related to immune response. Right panel. Significant GO categories exclusive to Alum+S vaccinated animals. (**D**) Plot depicting Log_2_ fold change of 28 identified genes (left) exhibiting differential expression patterns between the different conditions, and their associated cellular function (right). (**E**) Deregulation of selected cytokine genes including eosinophil chemo-attractant *Ccl11* and Th2 cytokine mRNAs in lungs of differently vaccinated hamsters upon SARS-CoV-2 infection. Heat map represents normalized mRNA expression (reads per kilobase per million bases mapped, RPKM) of genes. Grey cells with a X represents no detectable mRNA expression (RPKM= 0).

To investigate this in more detail, we performed gene ontology (GO) enrichment analyses to determine the biological processes associated with SARS-COV-2-induced DEGs in the hamster lung (Fig. 2C). For each group, enriched GO categories were associated with shared and distinct biological processes. In all groups, SARS-CoV-2 infection was associated with significant enrichment of genes associated with defense against pathogens, with vigorous induction of classical antiviral and inflammatory response genes (Fig. 2C, left panel). Of note, the magnitude of these responses was similar in unvaccinated and Alum+S immunized hamsters, and reduced in MeV_vac2_-SARS2-S(H) immunized animals. Visualizing significantly enriched GO categories that were unique to Alum+S treated animals revealed a profile of dysregulated gene classes that likely contributes to the observed vaccine induced pathology observed upon infection (Fig. 2C, right panel).

We next visualized fold change in expression of selected genes which displayed differential induction patterns between the groups, dividing the genes according to their associated cellular functions (Fig. 2D). These analyses led us to further explore normalized expression of a refined subset of genes which exhibited induction profiles unique to Alum+S immunization and which likely contribute to the vaccine-associated immunopathogenesis we observed (Fig. 2E). While the suite of genes involved eosinophil chemotaxis were similarly upregulated under all infection and vaccination conditions, uncontrolled induction of the major eosinophil chemotaxin *Ccl11* (eotaxin-1) mRNA was unique to hamsters vaccinated by alum-adjuvanted S protein (Fig. 2E). Furthermore, T_H_2 cytokine mRNAs *Il4*, *Il5*, *Il13* and *Il19* were potently induced in the majority of Alum-S vaccinated animals, but largely undetectable in the other groups. Taken together, these data describe the lung transcriptional signatures associated with MeV_vac2_-SARS2-S(H) induced protection in hamsters (Hörner et al. 2020). In parallel, these analyses also pinpoint likely transcriptional mediators underlying VAERD and hyperinflammation after T_H_2-biased vaccination: The specific induction of the *Il4*/*Il5*/*Il13*/*Il19* cytokine axis combined with the potent esosinophil chemotaxin *Ccl11* potentially results in uncontrolled recruitment of eosinophils to the site of infection.

### Induction of T_H_2-biased anti-S immunity in Alum-S hamsters

To further demonstrate induction of T_H_2-biased immunity as the trigger of the immunopathogenesis after SARS-CoV-2 challenge, Syrian hamsters were vaccinated as before with alum-adjuvanted S protein, MeV_vac2_-SARS2-S(H), or medium (MOCK). We then analysed antigen-specific immune responses of these animals without SARS-CoV-2 challenge. For this purpose, splenocytes of immunized hamsters were analysed 14 days after the second immunization in re-call experiments for antigen-specific induction of T_H_2 cytokines IL-4, IL-5, or IL-13.

To control successful vaccination of all cohorts first, IFN-γ secretion was determined by Enzyme-linked Immunosorbent Spot assay (ELISpot) after re-stimulation. Splenocytes of all animals reacted with IFN-γ secretion after stimulation with the unspecific stimuli ConA or Flagellin, which triggered more than 750 spots or 600 spots / 10^6^ cells, respectively, thus demonstrating functionality of all hamsters’ splenocytes. On the other hand, medium- or Ovalbumin-stimulated splenocytes showed a background reactivity of 100 to 250 IFN-γ^+^ spots / 10^6^ cells (Fig. 3A). Upon antigen-specific stimulation with recombinant S protein, only animals vaccinated with MeV_vac2_-SARS2-S(H) or alum-adjuvanted S revealed specific reactivity with approximately 400 spots / 10^6^ cells or in the range of the upper limit of detection, respectively, thereby demonstrating successful induction of cellular immune responses against the SARS-CoV-2 Spike protein by both vaccines.

**Fig.3:**
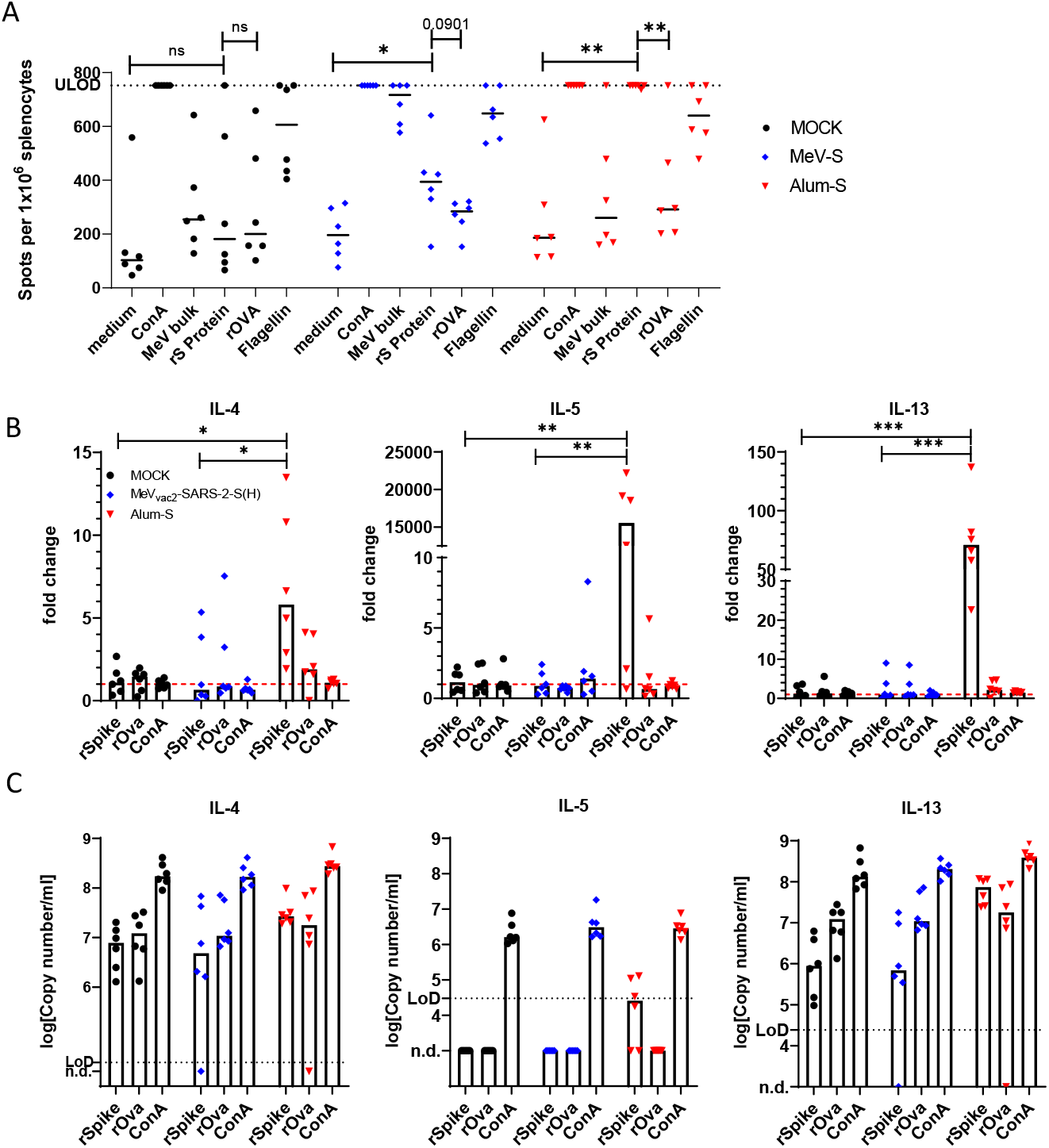
Induction of T_H_2-biased antigen-specific immune cells in protein-vaccinated Syrian hamsters. **(A)** IFN-γ ELISpot analysis using splenocytes of hamsters vaccinated on days 0 and 21, isolated 14 days after the boost immunization and after stimulation with recombinant S protein (rS protein) or MeV bulk antigens (MeV bulk). The reactivity of splenocytes was confirmed by Concanavalin A (ConA) [5 μg/mL], or Flagellin treatment [20 μg/mL]. Recombinant ovalbumin (rOva) or medium served as negative controls. The number of cells per 1 x 10^6^ splenocytes represent the amount of cells expressing IFN-γ upon re-stimulation. Dots represents individual animals, horizontals bars median per group (n = 6). Samples above the upper limit of detection (ULOD) were displayed as such. For statistical analysis of grouped ELISpot data, paired t test was applied. ns, not significant (p>0.05); *, p<0.05; **, p<0.01. **(B)** Relative fold-change expression of mRNAs encoding IL-4, IL-5, or IL-13 was determined using quantitative RT-PCR and the ΔΔct method. mRNA encoding RPL18 was used as housekeeping gene for normalization. Mean of samples from mock-treated hamsters served as reference. For statistical analysis, ordinary one-way ANOVA was applied with Tukey‘s multiple comparisons test. ns, not significant (p>0.05), *, p<0.05; **, p<0.01; ***, p<0.001. **(C)** Absolute mRNA copy numbers were determined by quantitative RT-PCR using a plasmid DNA standard for each analyte. Mock-vaccinated hamsters, black circles; MeV_vac2_-SARS2-S(H)-vaccinated hamsters, blue diamonds; protein-vaccinated hamsters, red triangles.

In the absence of suitable assays to determine hamster IL-4, IL-5, or IL-13 protein secreted by immune cells, we determined mRNA copy numbers of these cytokines by quantitative RT-PCR (qRT-PCR) in re-stimulated splenocytes and correlated the signals to the hamster housekeeping gene *RPL18* mRNA copies. For this purpose, the total RNA was prepared from splenocyte cultures after re-stimulation and subjected to analysis (Fig. 3B). For normalization of the signals, all were correlated to the average signal of splenocytes of mock-immunized hamsters after incubation with the respective stimuli (Fig. 3C). As expected, S-specific induction of *Il4* (5-fold), *Il5* (>15,000-fold), or *Il13* (70-fold) mRNA was only found in splenocytes of animals immunized by alum-adjuvanted S while splenocytes of MeV-vaccinated animals did not show any up-regulation of these T_H_2-cytokine genes after re-stimulation.

These data demonstrate that the MeV-derived vaccine did not induce S-specific T_H_2-biased immune cells after vaccination, in contrast to alum-adjuvanted Spike protein, corroborating the hypothesis of VAERD induction after SARS-CoV-2 challenge specifically by the T_H_2-biased alum-adjuvanted S protein.

### Replication of VAERD in hamsters responding to dexamethasone treatment

To evaluate treatment options for VAERD and to further dissect the role of individual cell populations in infected hamster lungs during VAERD, the initial challenge experiment of vaccinated hamsters was replicated in a third set of hamsters for the purpose of downstream scRNA-Seq analysis. For this purpose, cohorts of 4 to 6 Syrian hamsters were vaccinated as before with animals receiving alum-adjuvanted S, MeV_vac2_-SARS2-S(H), or medium (MOCK). For the cohort receiving Alum+S, the number of animals was doubled to be able to assess the impact of treatment with the clinically used immunosuppressive dexamethasone on VAERD.

To control the immunization and to be able to stratify the Alum+S vaccinated cohorts, sera of immunized animals were tested for binding antibodies targeting MeV or SARS-CoV-2 S by ELISA and neutralizing antibodies by titration of VNT (Suppl. Fig. S3). As expected, binding antibodies specific for MeV bulk antigens were detected solely in the post boost (day 31) sera of hamsters vaccinated with MeV_vac2_-SARS2-S(H). Consistently, binding antibodies to SARS-CoV-2 S were detected in sera from all animals that received adjuvanted protein or the MeV-derived vaccine (Suppl. Fig. S3B). However, neutralizing activity inhibiting SARS-CoV-2 infection was evident only in hamsters immunized with recombinant MeV_vac2_-SARS2-S(H) with a VNT of 10 to 80, but not in the protein-vaccinated animals. MeV neutralizing antibody (nAb) titers after boost immunization reached a VNT of 320 to 640 VNT (Suppl. Fig. S3H). The binding Ab titers targeting SARS-CoV-2 S were used together with the animals’ sex to stratify both cohorts immunized with the protein vaccine for treatment with dexamethasone upon challenge, or not. All animals were challenged 14 days after the second vaccination (i.e. on day 35) by intranasal inoculation of SARS-CoV-2 low-passage patient isolate MUC-1 (Hörner et al., 2020). During the next four days, one of both groups that had received the protein vaccine were treated twice daily with dexamethasone, the other was left untreated and monitored for disease progression.

Upon infection with SARS-CoV-2, weight loss was initially observed in all groups (Fig. 4A, Suppl. Fig. S4A). Consistent with previous observations, hamsters immunized with MeV_vac2_-SARS2-S(H) stopped losing weight by day 2 p.i., and recovered thereafter, while naïve animals revealed progressive weight loss over the course of the experiment. This was also evident for hamsters immunized with the protein vaccine that showed comparable decay, as observed before. Remarkably, the protein-vaccinated, infected animals treated with dexamethasone revealed another phenotype: weight loss stopped on day 2.5 and the animals’ weights stabilized for the next 2 days. All animals were sacrificed on day 4 p.i. and subsequently dissected.

**Fig. 4:**
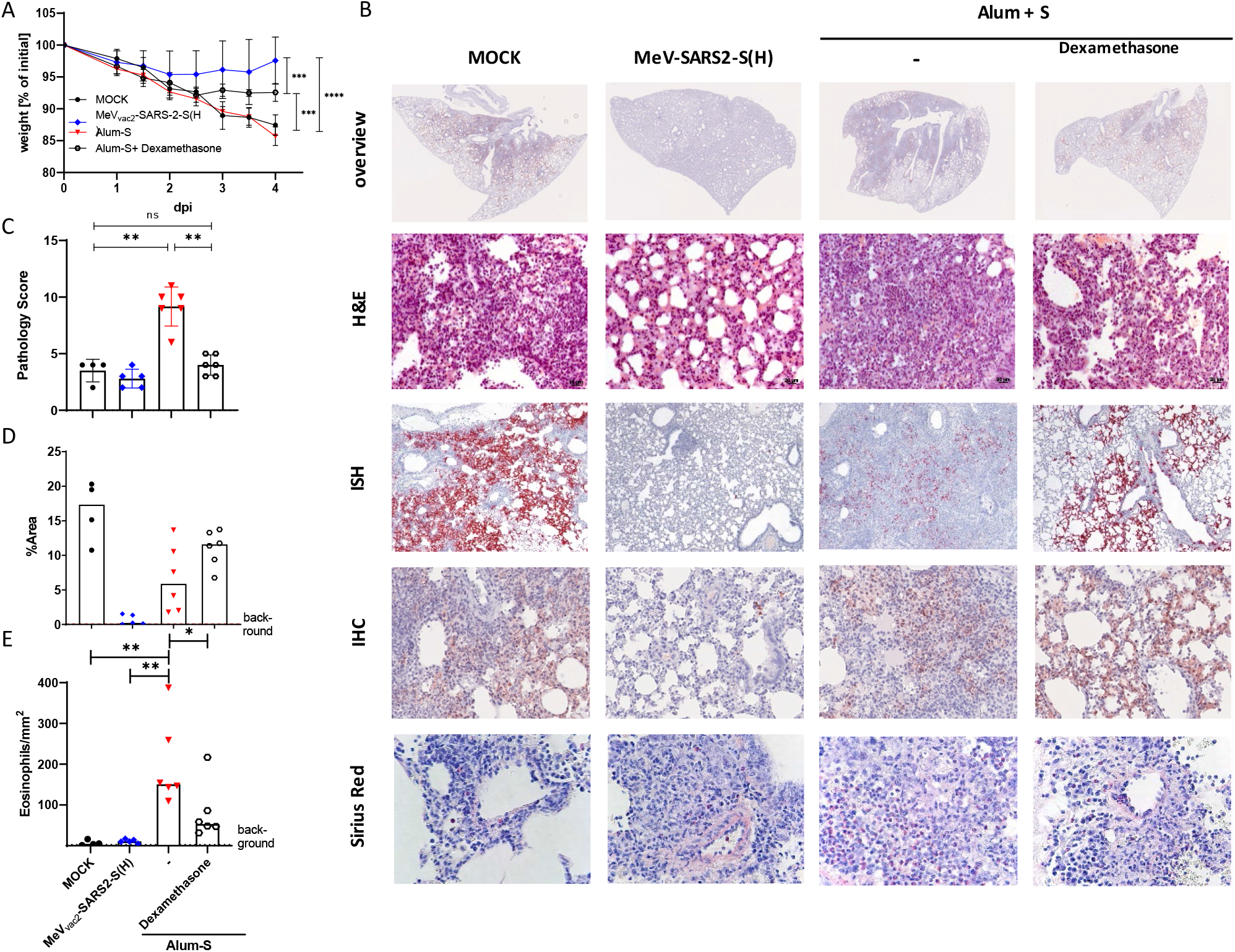
Protection of vaccinated hamsters and pathology upon SARS-CoV-2 challenge in combination with dexamethasone treatment. Immunized hamsters were intranasally challenged with low-passage SARS-CoV-2. **(A, B)** Protective efficacy was analysed by **(A)** quantifying weight loss over time and **(B)** histopathology of lung sections prepared 4 dpi (top row, 4x magnification; all other rows, 400x magnification). **(C)** Blinded histopathological analysis of H&E-stained lung samples (B, second row) was performed and findings were scored. **(D)** Infected areas positive by *In situ* hybridisation against SARS-CoV-2 RNA (ISH; B, 3rd row) were semiquantitatively scored. Infection was confirmed by immunohistochemistry against SARS-CoV-2 nucleocapsid protein (IHC; B, 4^th^ row). **(E)** Infiltration of eosinophils positive by sirius red staining (B, bottom row) was quantified by determination of the mean number of eosinophils per mm^2^ for each animal. Mock-immunized, black circles; MeV_vac2_-SARS2-S(H), blue diamonds; Alum+S without (red triangles) or with dexamethasone treatment (open circles).

Analysis of gross pathology replicated the pathology observed previously (Suppl. Tab. S2), with signs of pneumonia and inflamed areas on the lungs of naïve animals, while MeV_vac2_-SARS2-S(H) immunized hamsters showed few lesions on the lungs’ surface (Suppl. Fig. S4B). Moreover, this experiment replicated the VAERD observed before, which became evident already in the gross pathology, with large areas of inflammation on the surface of the S-protein vaccinated animals’ lungs and a swollen phenotype of the entire lung. This effect was prevented in the dexamethasone-treated animals. Their lungs revealed a largely normalized appearance of the lung explants comparable to the MeV_vac2_-SARS2-S(H) treated cohort. All animals’ lungs were subjected to broncheoalveolar lavage (BAL) and subsequently sampled, with specific sections of each lung prepared for histology (left lobe), analysis of total RNA (right middle lobe), titration of live virus (right apical lobe), and analysis of transcriptomics by scRNA-Seq (caudal lobe).

Histopathologic analyses of lung samples after H&E, ISH, immunohistochemistry and sirius red staining exactly replicated the VAERD phenotype for Alum+S vaccinated animals and absence thereof for MeV_vac2_-SARS2-S(H) vaccinated animals as described above for the first challenge experiment (Fig. 4B). When Alum+S immunized animals were treated with dexamethasone after challenge, the VAERD phenotype vanished also in the tissue pathology. Here, the inflammatory phenotype was absent and lung tissue appearance reflected the normalized phenotype of MeV_vac2_-SARS2-S(H) immunized hamsters. The pathology score decreased significantly under dexamethasone treatment and reflected naïve challenged animals (Fig. 4C). Despite this normalization of lung pathology, immunohistochemistry for SARS-CoV-2 N or ISH for SARS-CoV-2 genomes was not markedly different from naïve challenged animals and enhanced when compared to Alum+S vaccinated, untreated hamsters (Fig. 4B, last column).

Furthermore, live virus titers and viral RNA copy numbers, as determined in lung tissue (Fig. 5A, B) and bronchoalveolar lavage (BAL) cells (Fig. 5C), were in agreement with this phenotype. Virus RNA copy numbers within the lung tissue were significantly reduced in hamsters vaccinated with MeV_vac2_-SARS2-CoV (0.36 – 8.28 E-gene copies/RPL18 copy) compared to naïve infected animals (110.8 – 235.3 E-gene copies/RPL18 copy), consistent with the absence of live virus in the lungs of all animals in the MeV-group (Fig. 5A). Compared to naive/unvaccinated infected hamsters, viral burden was lower in hamsters immunized with Alum+S, but increased slightly when these hamsters were treated with dexamethasone during the challenge. Comparable virus RNA copy numbers were obtained for the BAL cells studied.

**Fig. 5:**
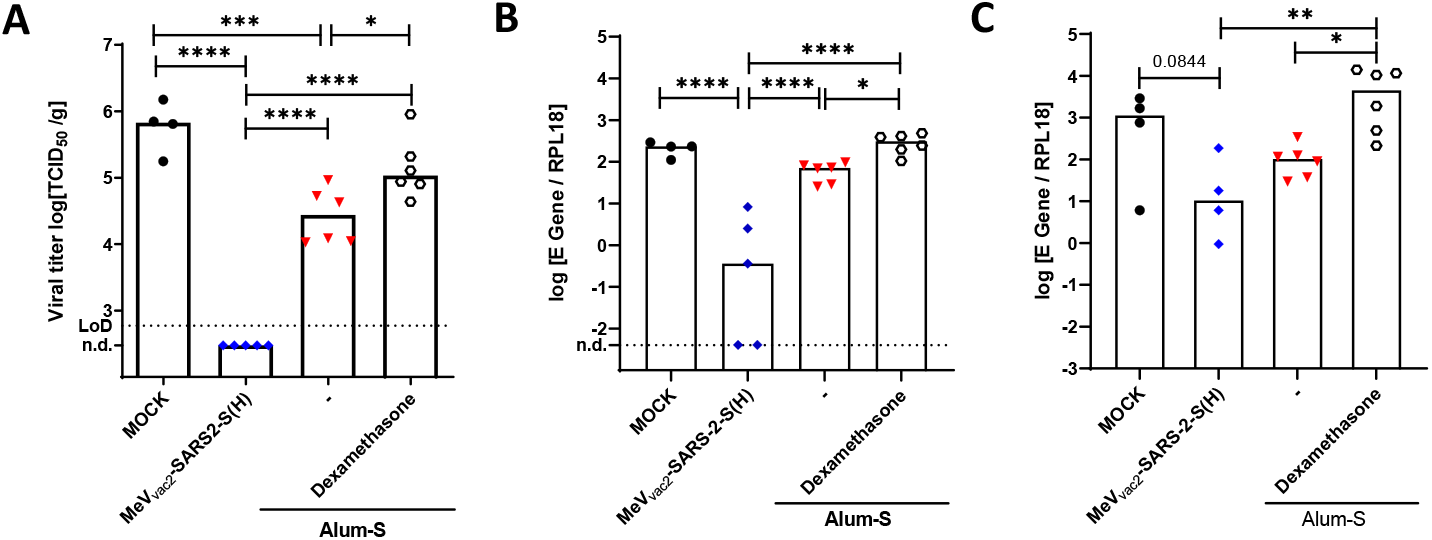
Impact of vaccination or dexamethasone treatment on SARS-CoV-2 titers upon challenge of Syrian hamsters. Protection of vaccinated hamsters upon challenge with SARS-CoV-2 was analysed 4 dpi by **(A)** titration or live virus titers in lung tissue, or **(B, C)** determination of relative SARS-CoV-2 E gene copy numbers in **(B)** lung tissue or **(C)** bronchoaveolar lavage (BAL) cells by quantitative RT-PCR. RPL18 housekeeping gene served for normalization. Lower limits of detection (LoD) are indicated by dotted lines. Each dot represents an individual animal vaccinated with medium (black circles), MeV_vac2_-SARS2-S(H) (blue diamonds), or Alum-adjuvanted Spike protein without (red triangles) or after treatment with dexamethasone (open circles). For statistical analysis, ordinary one-way ANOVA was applied with Tukey‘s multiple comparisons test. ns, not significant (p>0.05), *, p<0.05; **, p<0.01;***, p<0.001;****, p<0.0001.

Thus, viral loads and extent of tissue infection did not correlate with the inflammatory phenotype of pathology, consistent with an immunopathogenesis as the basis for VAERD, which again was not observed for the T_H_1-biased MeV-COVID-19 model vaccine candidate, but evident in all animals that received adjuvanted protein. Moreover, also the responsiveness of the pathology to dexamethasone treatment indicates implication of immune cells in this process.

### Dysregulation of T_H_2 cytokines in lungs and BAL cells

To confirm pulmonary up-regulation of *Il4*, *Il5*, *Il13*, and *eotaxin-1* mRNAs in protein-vaccinated animals and absence thereof in MeV-vaccinated animals and to determine if this dysregulation was also observed in BAL cells after infection, total RNA of the respective cells was subjected to qRT-PCR. All cytokine-encoding genes were assessed by the ΔΔct method normalized to the mean of naïve infected control animals. These analyses revealed significant induction of all four cytokine genes in lung cells and revealed the same tendency in BAL cells after the infection of protein-vaccinated animals. In contrast, vaccination with the T_H_1-biased measles-derived candidate down-regulated expression of these critical genes, while dexamethasone treatment did not alter the cytokine expression pattern (Suppl. Fig. S5). Therefore, induction of T_H_2 cytokines in protein-vaccinated animals identified by RNA-Seq in the first experiment was confirmed in this set of hamsters by analysis of individual mRNA populations.

### Assignment of dysregulation to individual cells in lungs by scRNA-Seq

To delineate the role of individual cell populations, scRNA-Seq data of infected lungs were generated and first analysed to determine cellular subsets according to individual cellular gene expression profiles, as described previously (Nouailles et al., 2021). By these means, it became possible to define 25 individual cell clusters (Suppl. Fig. S6), the transcriptional profile of which could be assigned to 13 different cell populations including immune cells (Fig. 6B). Minor differences between vaccine cohorts became evident in the relative cell frequencies in the infected lungs. In naïve animals and even more in Alum+S vaccinated animals, lung macrophages, that revealed traits of an interstitial macrophage phenotype, were overrepresented in comparison to the MeV-vaccinated group. In contrast, alveolar macrophages were observed at higher frequencies in MeV-vaccinated animals’ lungs (Fig. 6C).

**Fig. 6:**
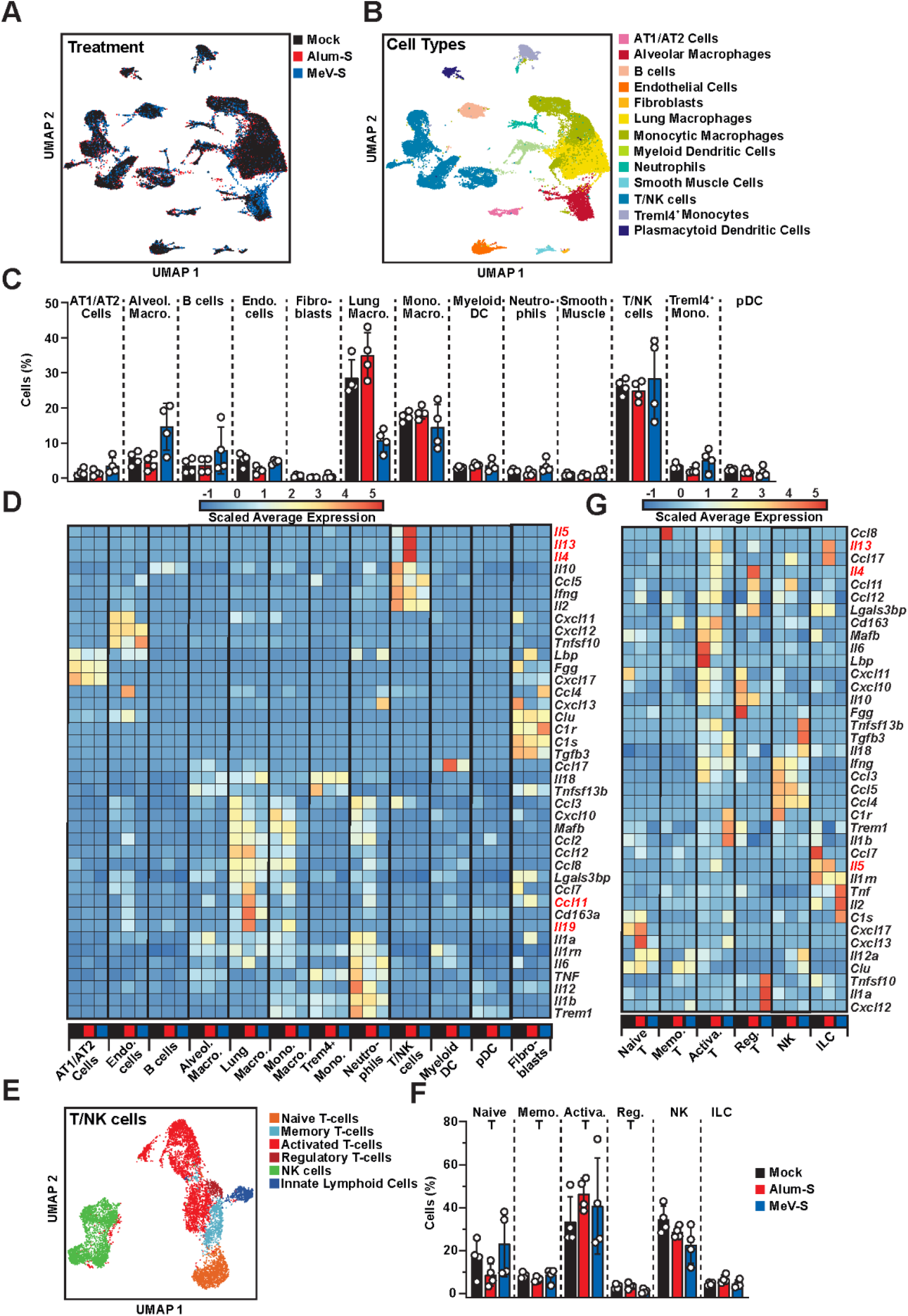
Annotation of cell populations in infected hamster lungs and respective gene regulation. **(A)** Definition, **(B)** annotation, and **(C)** quantification of specific cell populations found in the lungs of differently vaccinated hamsters 4 dpi. **(D)** Differently regulated genes of interest in these cell populations as indicated displayed by heatmap. To further pinpoint dysregulation of T or NK cells, the respective cell subsets were **(E)** annotated according to maker genes, **(F)** quantified and **(G)** de-regulation of genes was resolved in the respective heatmap as displayed. **(A, C, D, F, G)** Black, mock-vaccinated hamsters; red, hamsters vaccinated with Alum-adjuvanted Spike protein (Alum+S); blue, hamsters vaccinated with MeV_vac2_-SARS2-S(H) (MeV-S).

Analysing the gene expression profiles of these distinct cell populations, specific expression patterns became evident (Fig. 6D). Up-regulation of CCL-11 expression observed exclusively in the Alum+S group could be assigned to lung macrophages, which were overrepresented in this cohort (Fig. 6D). Induction of *Il4*, *Il5*, and *Il13* became evident exclusively in the population of T- and NK-cells, in accordance with the results of the vaccination experiments (Fig. 6D, three top panels). Zooming in on specific T/NK cell subsets (Fig. 6 E, F), up-regulation of *Il4* was found in regulatory T cells, while *Il5* and *Il13* expression was assigned to cells with an innate lymphoid cell phenotype (Fig. 6G). In contrast, MeV_vac2_-SARS2-S(H) vaccinated animals reflected a similar, but dampened response when compared to naïve infected animals, with few genes being differentially regulated. On the other hand, monitoring the distribution of SARS-CoV-2 RNA sequences across the different cell types as a measure for virus infection or uptake, a broad presence of viral RNA in most cell types was observed in naïve/unvaccinated, infected animals (Fig. 7A, B). This distribution was more focused in lung macrophages in Alum+S vaccinated animals. This correlated to some extent with the expression profiles of Fcγ-receptors IIb and IV, which were found to be up-regulated after infection in naïve and Alum+S vaccinated samples. In particular Fcγ-R IIb showed remarkable up-regulation in the Alum+S cohort, when compared to naïve animals (Fig. 7C).

**Fig. 7:**
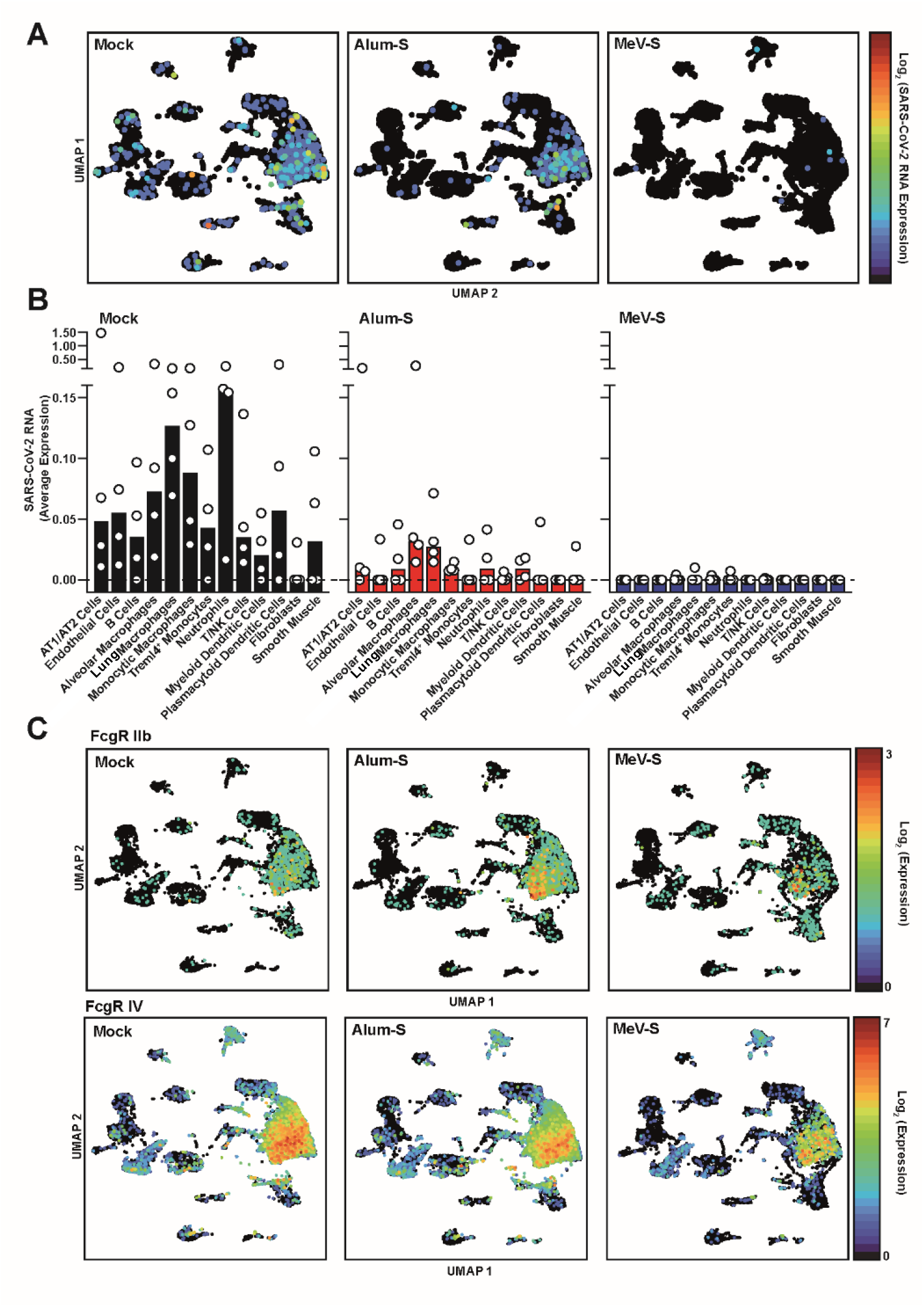
Assignment of SARS-CoV-2 reads and FcR-expression to cell populations in infected hamster lungs. **(A)** Assignment of SARS-CoV-2 genome reads to cell populations defined in Fig. 6 and **(B)** quantification of respective positive cells of the different treatment groups as indicated. Single dots represent individual animals. **(C)** Assignment of mRNA reads encoding hamster FcγR IIb (upper panel) or FcγR IV (lowerpanel) to cell populations defined in Fig. 6.

To further mine the scRNA-seq data, we performed both gene set variation analysis (GSVA) and gene set enrichment analysis (GSEA) using the REACTOME, KEGG and GO databases on specific cell populations which contribute to the VEARD observed in Alum+S vaccinated animals (Suppl. Fig. S7 to S9), focusing on alveolar macrophages, lung macrophages, monocytic macrophages, *Treml4^+^* monocytes, and T/NK cells. For GSVA, normalized, by-cell-population gene expression values were used as input, while DEGs between the unvaccinated and either Alum+S or MeVvac2-SARS2-S(H) vaccinated conditions respectively were utilised for GSEA. These analyses highlight pathways and biological processes, which are induced by SARS-CoV-2 infection and are differentially targeted or regulated across these selected cell types, dependent on prior immune status and vaccination type. These supportive analyses of the hamster lung scRNA-seq data provides insights into how the observed transcriptional differences mediate their downstream effects, exhibiting differential targeting of a broad range of cellular process or canonical pathways in a cell-type specific manner.

Taken together, the scRNA-Seq data assigned the up-regulation of IL-4, IL-5, and IL-13 to specific T-cell subpopulations induced by vaccination with alum-adjuvanted Spike glycoprotein, while CCL-11 expression was contributed by lung macrophages, which were overrepresented, revealed up-regulation of Fcγ-receptors and were the main target population containing an excess of SARS-CoV-2 RNA. These aberrant patterns of dysregulated gene expression were not observed in animals vaccinated with the prototypic T_H_1-biased MeV-COVID-19 vaccine candidate.

## Discussion

Based on these data, we propose the following mechanism for induction of VAERD in our animal model (Fig. 8): Vaccination with alum-adjuvanted S protein in post-fusion conformation induces low, non-protective levels of S-specific binding antibodies lacking neutralizing activity. In parallel, T_H_2-biased S-specific T cell responses were induced as evident by the significant up-regulation of IL-4, IL-5, and IL-13 after recall. After infection, these immune responses lower the virus load to some extent, but the induction of T_H_2-immunity ends up in VAERD via the IL-4/IL-5/IL-13 chemokine axis secreted by regulatory T cells and innate lymphoid cells. These processes trigger attraction of eosinophils via chemoattractants such as CCL-11 secreted by hyperstimulated lung macrophages that enrich SARS-CoV-2 virus particles and are most likely stimulated by these particles potentially via Fcγ-receptor mediated up-take of opsonized viruses as proposed for antibody-dependent enhancement (ADE) processes.

**Fig. 8:**
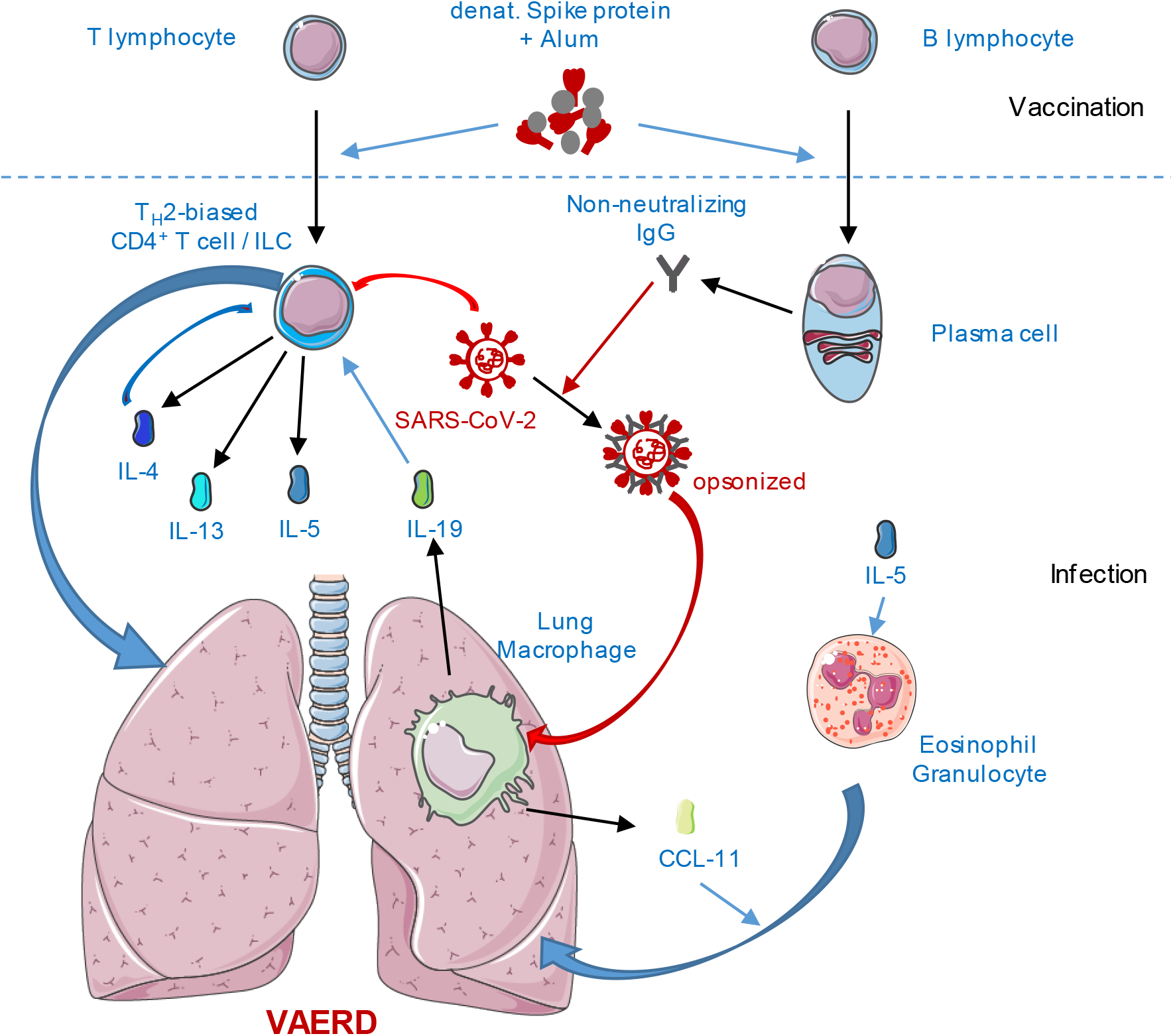
Schematic depiction of VAERD induction by T_H_2-biased COVID-19 vaccines. After vaccination with alum-adjuvanted (grey) Spike protein (red), T_H_2-biased, S-specific CD4+ T helper cells are induced and B cells are stimulated to secrete non-neutralizing antibodies (black) (Upper part of scheme). After infection with SARS-CoV-2 (red) virus stimulates S-specific T cells to produce IL-4, IL-5 and IL-13 (blue), setting the stage for lung disease. In parallel, virus particles are opsonized by non-neutralizing antibodies and are taken up by interstitial macrophages that subsequently secrete CCL-11 (light green) that attracts eosinophils, which become also activated by IL-5. In parallel, macrophages secrete IL-19 (green) that re-enforces T_H_2-polarization of the T cell responses, as does IL-4 in a positive feedback loop (lower part of the scheme). Altogether, these processes orchestrate hyperinflammation of the lung, i.e. VAERD, comparable to allergic asthma. Assembled with material from Servier Medical Art by Servier(smart.servier.com).

While our data strongly support such a model for induction of VAERD by sub-optimal T_H_2-biased prototypic vaccines targeting SARS-CoV-2, the nature of these findings seems quite striking. VAERD had been clearly observed for both MERS-CoV and SARS-CoV in the respective transgenic mouse models after vaccination with T_H_2-biased whole-inactivated virus vaccines (Bolles et al., 2011; Tseng et al., 2012; Iwata-Yoshikawa et al., 2014; Honda-Okubo et al., 2015; Agrawal et al., 2016). Since no human CoV vaccines had been tested against these two pathogens, the transferability of these findings to the occurrence of VAERD in humans remained unclear. Nevertheless, this potential risk was perceived also for vaccine-induced immune responses that target the closely related SARS-CoV-2, and triggered the developers to reach for T_H_1-biased immunity and to assess the potential of immunopathogenesis in the available animal models (Anderson et al., 2020; Corbett et al., 2020b; Corbett et al., 2020a; Jackson et al., 2020; Polack et al., 2020; Ramasamy et al., 2020; Walsh et al., 2020a; Sadoff et al., 2021; Stephenson et al., 2021; van der Lubbe et al., 2021). Despite all these efforts, only one study has been published so far that revealed evidence for VAERD potential upon vaccination of T_H_2-prone Balb/c mice with wilfully denatured antigen followed by a challenge with a mouse-adapted recombinant SARS-CoV-2 (DiPiazza et al., 2021). No other studies so far have identified evidence for the risk of enhanced disease upon COVID-19 vaccination. How can this discrepancy be explained?

In previous studies, we primarily focused our efforts on the efficacy of our T_H_1-biased COVID-19 vaccine (Hörner et al., 2020), which indeed revealed, as a prototype T_H_1-biased vaccine, no evidence at all for immunopathogenesis. However, we included a sub-optimal vaccination regime by vaccinating Syrian hamster sub-cutaneously with Alum-adjuvanted, non-stabilized S protein expected to give a rather mediocre (Nürnberger et al., 2019), but T_H_2-biased immune response (Ko et al., 2017) as a worst-case control. Indeed, only binding antibodies with no neutralizing activity were induced, and no antigen-specific CD8^+^ T cell killing activity was observed either in this study, or before by us (Hörner et al., 2020) or others (DiPiazza et al., 2021). Thirdly, the hamster challenge model used in our experiments takes advantage of a couple of factors: A very low-passage primary patient isolate was used that did not reveal any tissue culture adaptation such as loss of the furin-cleavage site, which are known to impair pathogenicity of SARS-CoV-2. Inoculated into Syrian hamsters, this isolate causes a disease phenotype that closely mirrors moderate to severe forms of COVID-19, in contrast to other strains causing mild to moderate disease (Muñoz-Fontela et al., 2020).

This is very different from disease models using tissue-culture adapted virus in hamsters or other animal models. While transgenic K18-ACE2 are no doubt useful for studying protection by vaccines, antibodies or antivirals against an exaggerated generalized form of the disease (Winkler et al., 2020), the predictability of the CoV-receptor transgenic mouse model for disease course in human patients seems somewhat questionable, especially with regards to lethal neuroinvasion observed. On the other hand, non-human primates rarely develop a severe phenotype of disease after contact with SARS-CoV-2 (Shou et al., 2021).

Most likely, the combination of all these factors, i.e. a low, non-functional T_H_2-biased Ab responses potentially mimicking deteriorating immunity over time with an animal challenge model closely mimicking medium to severe COVID-19 allowed us to trigger and to detect VAERD in this very special setting. Our observations nevertheless closely resemble VAERD induced by inactivated RSV ((Johnson and Graham, 1999; Swart et al., 2002; Johnson et al., 2003; Johnson et al., 2004; Moghaddam et al., 2006) or MeV in the respective animal models (Polack et al., 2003) that are quite reminiscent of the situation observed in human VAERD after RSV (Openshaw, 2001) or measles vaccination (Nader and Warren, 1968) using whole-inactivated virus vaccines.

As already pointed out by DiPiazza et al, drawing conclusions from animal models and extending these observations to the immunological situation in humans may be a difficult task (DiPiazza et al., 2021), considering the limitations of the system, especially in the absence of experience with vaccines targeting SARS-CoV and MERS-CoV and evidence for VAERD of all vaccine concepts including whole-inactivated viruses in humans. However, we were able to replicate VAERD induced by a protein based COVID-19 vaccine candidate with a very similar phenotype in a second animal model, Syrian hamsters, using an unmodified low-passage virus isolate. Therefore, our data strongly support the idea to monitor vaccinated human patients that experience a break-through infection closely. In any case, while our experimental vaccines mimic, but are not the same as the authorized vaccines, these and previously published data by diPiazza et al. point at considerably few concerns for vaccines developed to trigger T_H_1-biased responses such as viral vector platform-based vaccines and mRNA vaccines. Moreover, even if VAERD as observed in our model should occur in human patients, this immunopathology would be treatable by dexamethasone, which revealed to be an effective medication for severe courses of COVID-19, anyway (Tomazini et al., 2020; Horby et al., 2021). This would be good news also for putative VAERD being mistakenly diagnosed as a variant of the usual forms of severe COVID-19 in a naïve patient.

Finally, our data indicate that ADE-like processes as reported by Wan et al. for MERS-CoV could be relevant in the mechanism of VAERD. For MERS-CoV, a monoclonal Ab binding to the RBD of the S is able to cross-link MERS-CoV S and Fc receptors. Tested in a pseudovirus assay, this monoclonal Ab mediated virus entry into D32A-expressing cells and macrophages (Wan et al., 2020). Such an uptake mechanism would explain the enrichment of SARS-CoV-2 genomes in macrophage populations correlating with the Fc-receptor distribution as observed in our study. Enrichment and stimulation of this immune cell population result in the secretion of the major eosinophil attractant CCL-11 and IL-19, which will drive T_reg_s into T_H_2-polarization. Together with the IL-4/IL-5/IL-13 chemokine axis of such S-specific T_H_2-biased CD4 helper cell populations, CCL-11 will cause infiltration of eosinophils in a process mechanistically reminiscent of allergic asthma and therefore end up in the immunopathogenesis observed in this study.

## Acknowledgements

This work was supported by grants of the German Center for Infection Research (DZIF; TTU 01.805, TTU 01.922_00) and the German Ministry of Health (CHARIS) to M.D.M. and by funding of Berlin Institute of Health (BIH) to C.G.. J.K. is supported by the Center of Infection Biology and Immunity (ZIBI) and Charité PhD Program. G.N. is supported by the BMBF and by the Agence nationale de la recherche (ANR) in the framework of MAPVAP (16GW0247). The authors would like to thank Daniela Müller, Mona Lange, and Silvia Schuparis for excellent technical assistance, Bevan Sawatsky for support of BSL-3 procedures, Elke Völker for assistance with CD spectroscopy, the team of the animal husbandry for logistic support with hamster experiments, and Christoph Schürmann for scientific discussions. The authors are indebted to Klaus Cichutek for intramural funding and support, and to Stefan Schülke for providing recombinant ovalbumin and flagellin A. The SARS-CoV-2 Trimeric Spike (Cat.No. 101007) was obtained from the National Institute for Biological Standards and control, UK. Our thanks go to Dr. Barnes Graham, NIAID and Chris Ball, NIBSC for providing this protein.

## Author contributions

CH, GN, RJPB, and MDM conceptualized the study; DT, AG, DP, and RJPB took care of data curation; MDM and RJPB acquired funding to support the study; AE, SM, JK, AA, MA, PG, MN, RP, CM, MGS, AB, CK, EW, AK, and MDM performed experiments and collected data; DT, AG, DP, CM, EW, and RJPB analysed sequencing data; GN, CG, RJPB, and MDM managed and coordinated the project; RE contributed critical material for the study; ZI, ZW, SP, ML, GN, CG, RJPB, and MDM supervised the experiments and data analyses; AE, JK, DT, DP, EW, RJPB, and MDM prepared figures to visualize the data; AE, RJBP, and MDM wrote the original draft of the manuscript; all authors reviewed and edited the final manuscript.

## Data availability

RNA-seq and scRNA-seq data generated in this study and were submitted to the NCBI GEO database under accession number XXXYYYY. Any code used in the analysis of the data has been deposited at https://github.com/GoffinetLab/Ebenig_SARS-CoV-2_TH1-vs-TH2-Vaccines

