## Supplementary material for "In contrast to T_H_2-biased approaches, T_H_1 COVID-19 vaccines protect Syrian hamsters from severe disease in the absence of dexamethasone-treatable vaccine-associated enhanced respiratory pathology": Suppl. Fig. S1 - S9; Suppl. Tab. S1 - S3

### Ebenig et al. – Suppl. Figure S1

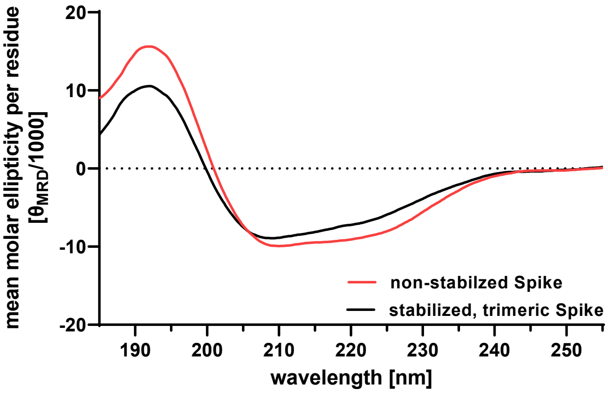


**Suppl. Fig. S1: Relative secondary structural elements of S determined by circular dichroism (CD).** To determine misfolding of recombinant Spike used for vaccination, spectra of non-stabilized recombinant S used for vaccination (red) and soluble S stabilized in the native conformation (black), secondary structure of the proteins was determined by CD spectroscopy. CD spectra of both proteins were recorded at room temperature from 255 nm – 185 nm by accumulating 10 runs, blotted and analysed for relative content of $\alpha$-helical and β-sheet secondary structural elements.

**Ebenig et al. – Suppl. Figure S2**

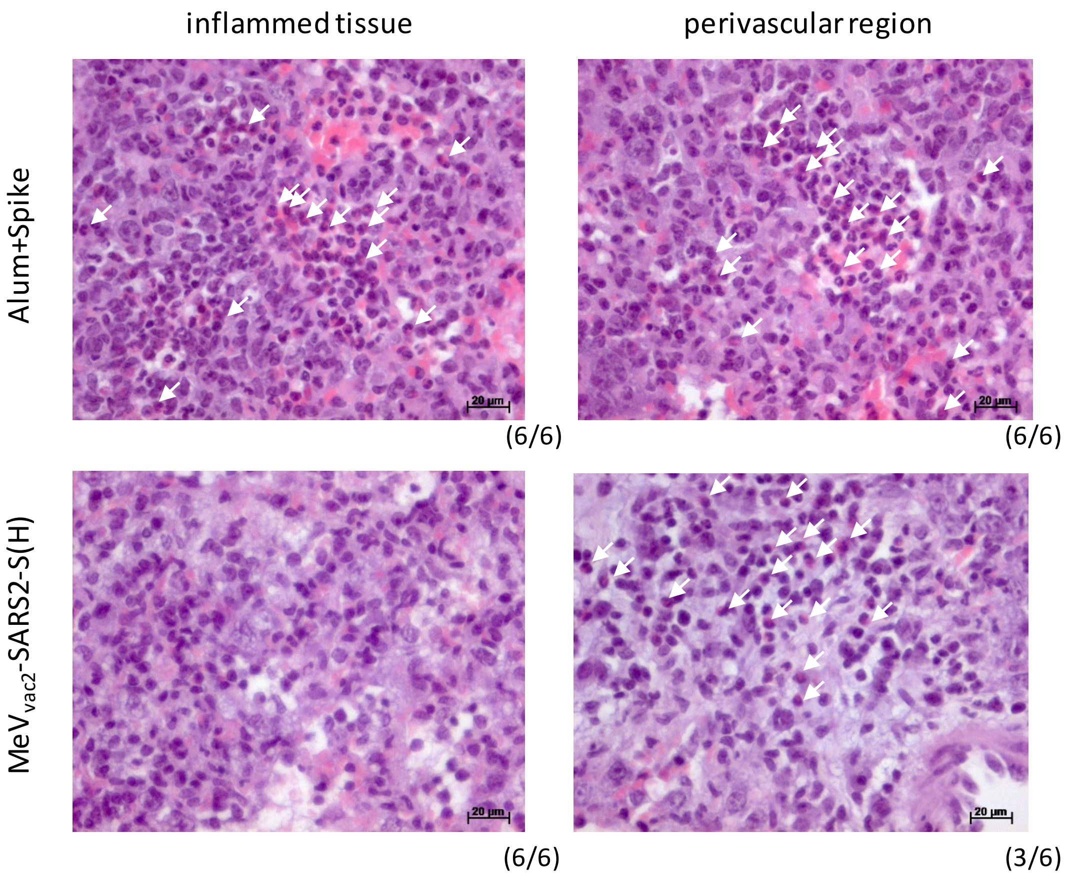


**Suppl. Fig. S2: Eosinophil infiltration into tissue of infected hamster lungs.** Haematoxilin and Eosin-staining of fixed lung slices of hamsters infected with SARS-CoV-2 after vaccination with Aluminum-adjuvanted Spike protein (upper panel) or $MeV_{vac2}$-SARS2-S(H) (lower panel) revealed eosinophil infiltration into inflammed tissue (left panel) of Alum+S, but not $MeV_{vac2}$-SARS2-S(H) immunized animals´. While eosinophils became evident in the perivascular region of samples of all animals vaccinated with protein before infection, only half (3/6) of the MeV-vaccinated animals revealed this phenotype. N = 6; representative pictures for the fraction of animals indicated below each picture. White arrows depict single eosinophils. Scale bar, 20 µm.

### Ebenig et al. – Suppl. Figure S3

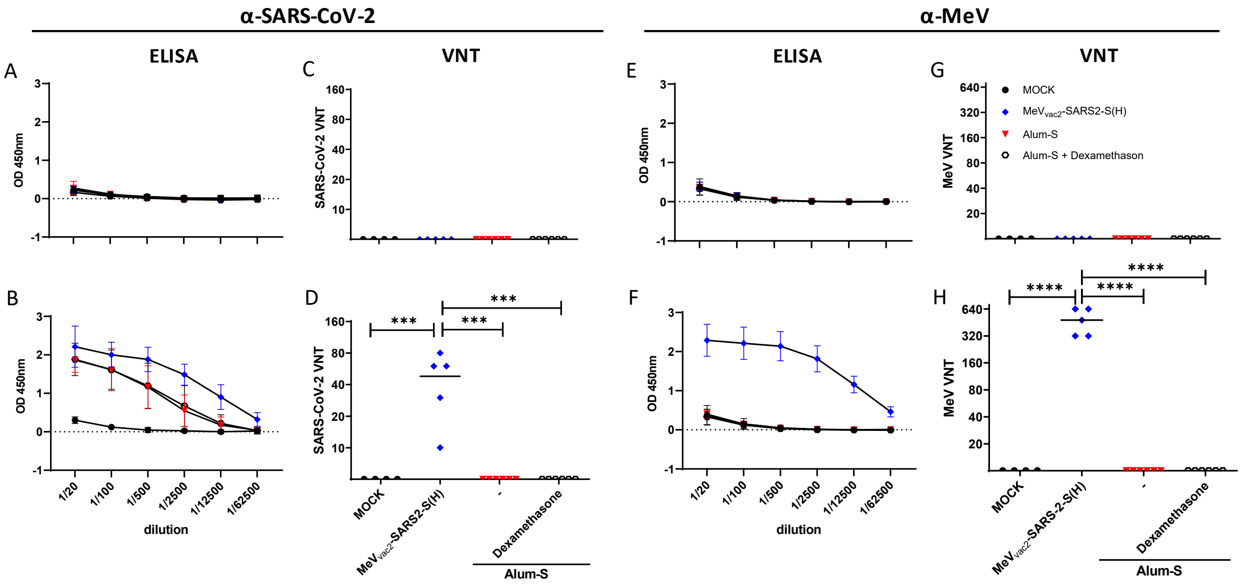


**Suppl. Fig. S3: Induction of α-SARS-CoV-2 S and α-MeV specific antibodies**. Sera of hamsters vaccinated on days 0 and 21 with $MeV_{vac2}$-SARS2-S(H) (blue diamonds) or Alum-adjuvanted S protein (red triangles, open circles) were collected on days 0 (A, C, E, G) and 31 (B, D, F, H) and analyzed for antibodies specific for SARS-CoV-2 S or MeV. Medium-inoculated hamsters (black circles) served as mock. Pan-IgG binding to recombinant SARS-CoV-2 S (A, B) or MeV bulk antigen (E, F) were determined by ELISA via the specific OD 450 nm value. Depicted are means and the respective standard deviation of each group (n = 4 - 6). Virus-neutralizing titers (VNT) in vaccinated hamsters for SARS-CoV-2 (C, D) or MeV (G, H) were calculated as the reciprocal of the highest dilution abolishing infectivity. For statistical analysis, ordinary one-way ANOVA was applied with Tukey's multiple comparisons test. ns, not significant (p>0.05), *, p<0.05; **, p<0.01;***, p<0.001;****, p<0.0001.

### Ebenig et al. – Suppl. Figure S4

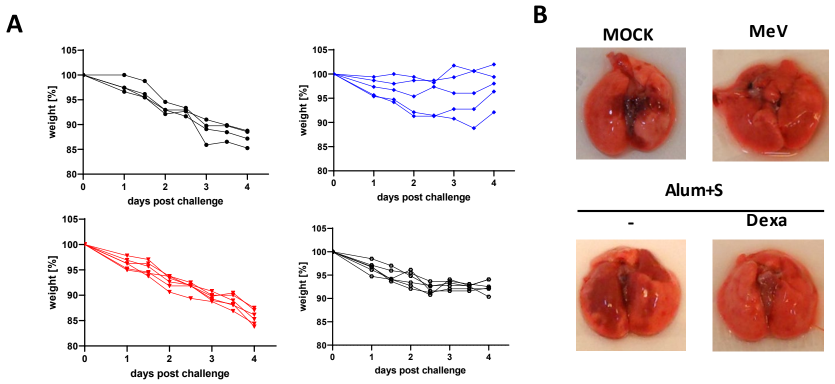


**Suppl. Fig. S4: Gross pathology in vaccinated hamsters after challenge.** Hamster were vaccinated at days 0 and 21 and challenged on day 35 with low-passage SARS-CoV-2. **(A)** Body weight changes of animals vaccinated with medium (upper left, black circles), $MeV_{vac2}$-SARS2-S(H) (upper right, blue diamonds), Alum+S without (lower left, red triangles) or with dexamethasone-treatment after challenge (lower right, open circles). **(B)** Macroscopic pathology of Syrian hamster lungs after SARS-CoV-2 infection and indicated vaccination or treatment on day 4 pi.

**Ebenig et al. – Suppl. Figure S5**

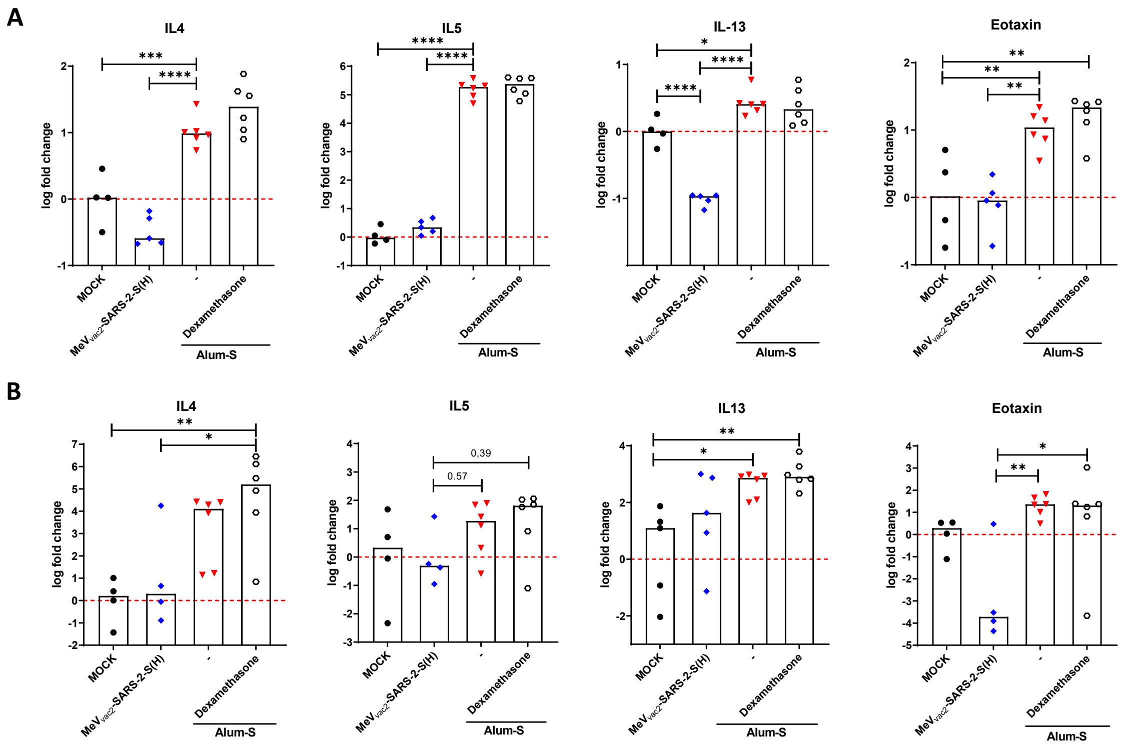


**Suppl. Fig. S5: Deregulation of $T_H2$ cytokines in lungs (A) and BAL cells (B) of SARS-CoV-2 infected vaccinated Syrian hamsters.** Relative fold-change expression of mRNAs encoding IL-4, IL-5, or IL-13 was determined using quantitative RT-PCR and the ΔΔct method. mRNA encoding RPL18 was used as housekeeping gene for normalization. Median of samples from mock-treated hamsters served as reference to normalize relative gene expression. Dots represent individual animals; mock-vaccinated hamsters, black circles; MeV$_{vac2}$-SARS2-S(H)-vaccinated hamsters, blue diamonds; protein-vaccinated hamsters, red triangles. For statistical analysis, ordinary one-way ANOVA was applied with Tukey's multiple comparisons test. ns, not significant ($p>0.05$), *, $p<0.05$; **, $p<0.01$;***, $p<0.001$;****, $p<0.0001$.

### Ebenig et al. – Suppl. Figure S6

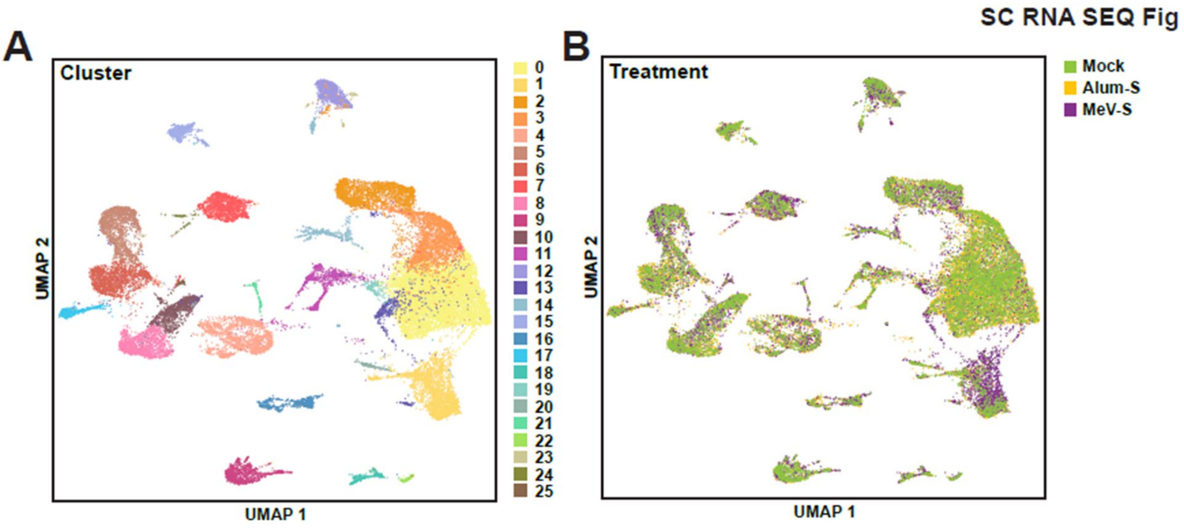


**Suppl. Fig. S6: Definition of cell populations in infected hamster lungs. (A)** Definition and **(B)** distribution of specific cell populations found in the lungs of vaccinated hamsters 4 dpi depending on the vaccination regime.

**Ebenig et al. – Suppl. Figure S7**

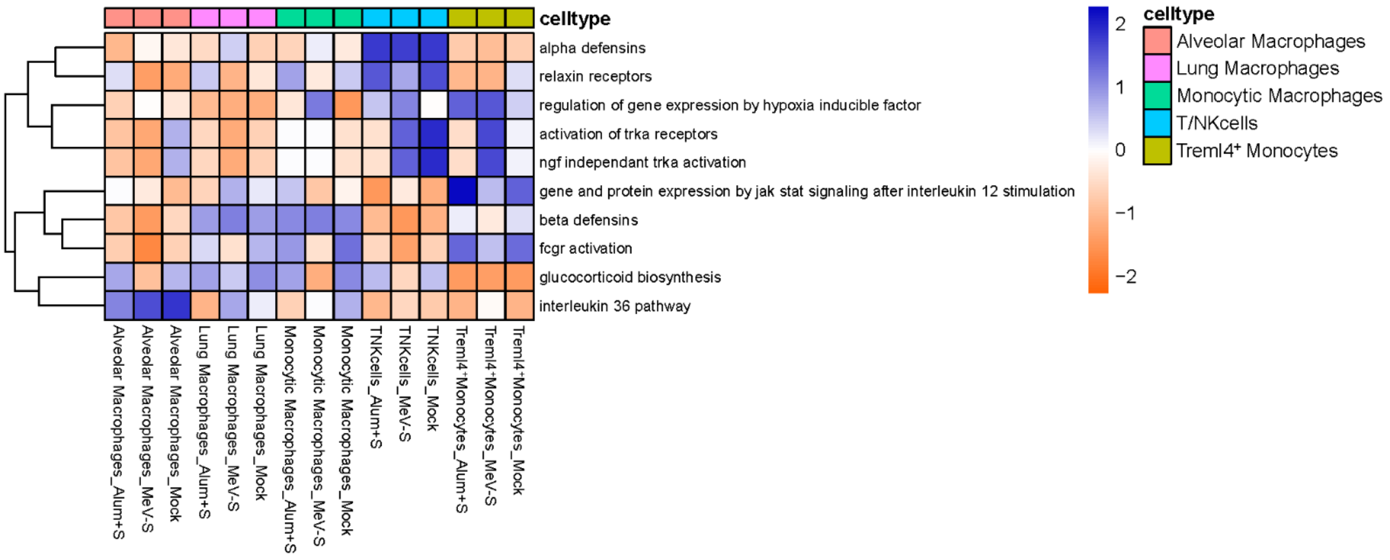


**Suppl. Fig. S7: SARS-CoV-2 infection triggers cell-type specific induction of REACTOME pathways which differ depending on vaccination type and prior immune status.** Heat map is derived from scRNA-seq data and highlights selected REACTOME pathways which exhibit differential activation or suppression triggered by infection. Differences are dependent on prior immune status or vaccination type, and are cell-type specific. Cell population specific activation or suppression of pathways observed in alveolar macrophages, lung macrophages, monocytic macrophages, Treml4$^{+}$ monocytes and T/NK cells is presented. Heat map is colored relative to the activation z-score gradient presented in the scale bar.

#### Ebenig et al. – Suppl. Figure S8

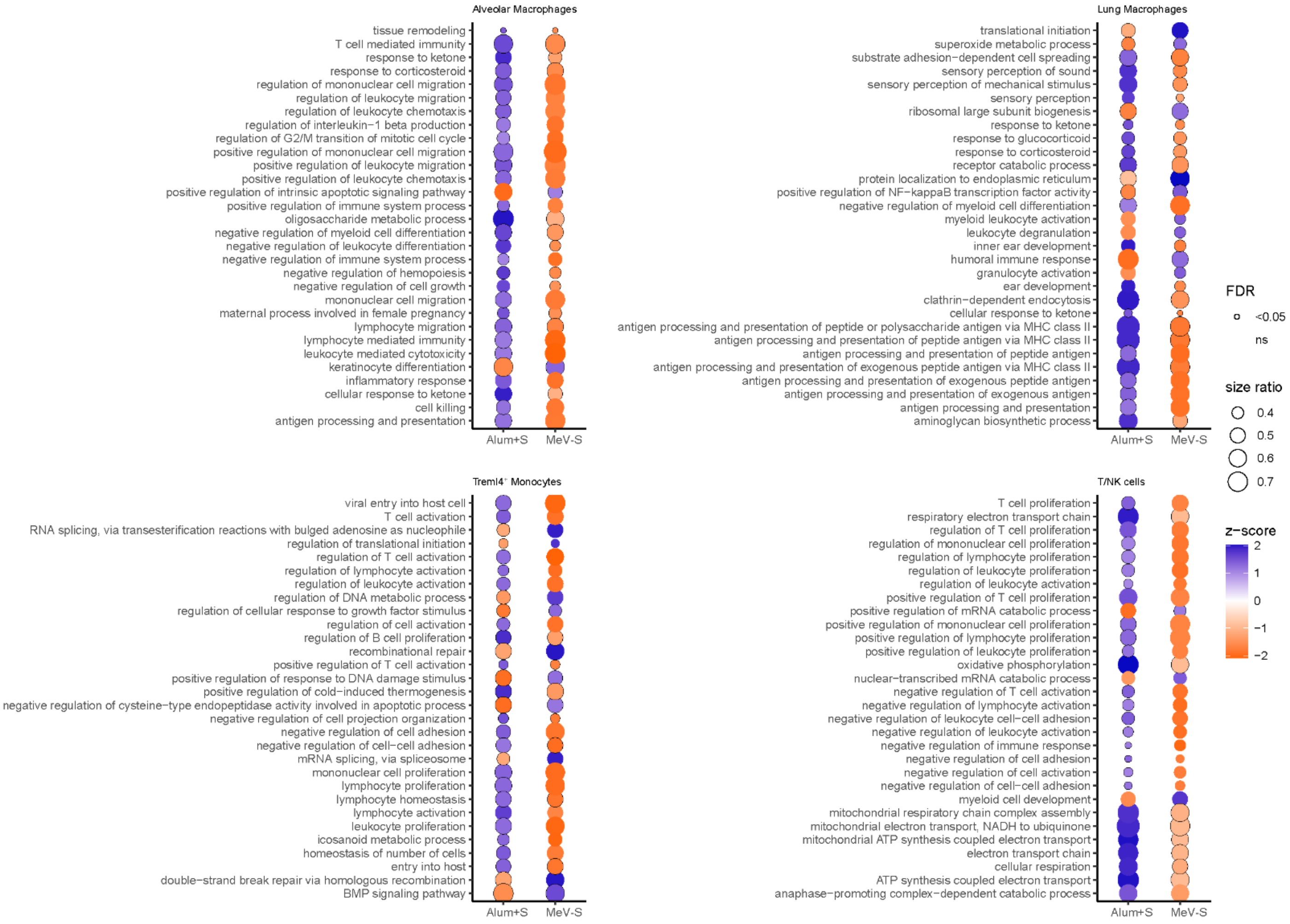


**Suppl. Fig. S8: SARS-CoV-2 infection induces cell-type specific enrichment of GO terms which differ dependent on prior vaccination type.** Dot plots highlight enriched GO categories that exhibit differential activation status dependent on prior vaccination with either Alum+S or $MeV_{vac2}$-SARS2-S(H) (MeV-S). Individual comparison plots are derived from scRNA-seq data and represent differential enrichment observed in alveolar macrophages, lung macrophages, Treml4$^{+}$ monocytes and TN/K cells. Enriched categories are labelled on the y-axes. Circle size represents the ratio of significantly dysregulated genes relative to the total gene number in a specific GO term. Circles are colored relative to activation z-score gradient presented in the scale bar, with significantly enriched GO categories (FDR p<0.05) highlighted with a black boarder. ns: non-significant

### Ebenig et al. – Suppl. Figure S9

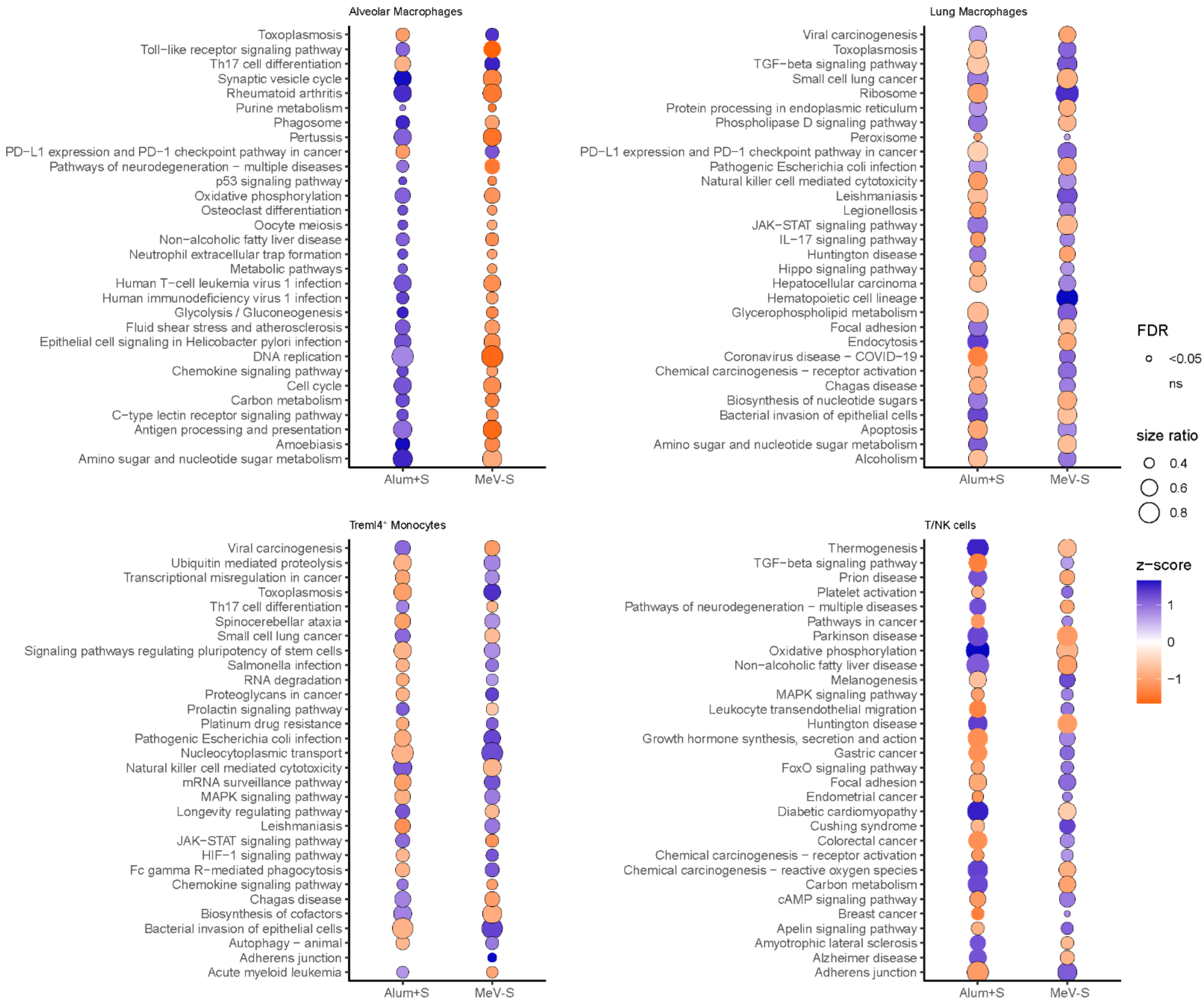


**Suppl. Fig. S9: SARS-CoV-2 infection induces cell-type specific activation or suppression of KEGG pathways which differ dependent on prior vaccination type.** Dot plots highlight targeted KEGG pathways that exhibit differential activation status upon SARS-CoV-2 infection, dependent on prior vaccination with either Alum+S or $MeV_{vac2}$-SARS2-S(H) (MeV-S). Individual comparison plots are derived from scRNA-seq data and represent differential enrichment observed in alveolar macrophages, lung macrophages, Treml4$^{+}$ monocytes and T/NK cells. Enriched pathways are labelled on the y-axes. Circle size represents the ratio of significantly dysregulated genes relative to the total gene number in a specific KEGG pathway. Circles are colored relative to activation z-score gradient presented in the scale bar, with significantly enriched KEGG pathways (FDR p<0.05) highlighted with a black boarder. ns: non-significant

**Supplementary Table 1. Histopathological analysis of lung tissue of vaccinated Syrian hamsters batch #1 upon challenge with SARS-CoV-2.** Hamsters were vaccinates with medium (MOCK), $MV_{vac2}$-ATU(P) as measles-only vector control (MeV), $MeV_{vac2}$-SARS2-S(H) (MeV-S), or Alum-adjuvanted Spike protein (Alum+S). The left lobe of vaccinated hamster lungs was dissected 4 dpi. H& E staining revealed histopathological changes and immune cell infiltration assessed by a trained pathologist in a blinded manner. DIC, disseminated intravascular coagulation.

| Animal-No. | Vaccine group | % Dense Area | Bronchia | Vessels | DIC | Dense Area | Fibrosis | Syncytia |
|---|---|---|---|---|---|---|---|---|
| 47 | MOCK | < 50% dense areas, clearly bronchial-associated | Marked purulent bronchitis (granulocytes and lymphocytes in the lumen),<br>bronchial epithelium sometimes with inflammatory infiltration. | Vascular walls with inflammatory infiltration,<br>Single-cell necrosis in the vessel wall,<br>multiple hemorrhages in the tissue,<br>2 x edema around vessels | No | mostly lymphocytes and macrophages, less with granulocytes, sporadic eosinophils, proliferation of type II pneumocytes, often karyorrhexis | No | No |
| 48 | MOCK | 30% dense areas, Bronchial- and vessel-associated | Bronchitis (lymphocytes and only very few granulocytes in the lumen), bronchial epithelium with little inflammatory infiltration and damage. | Vascular walls with inflammatory infiltration,<br>Single-cell necrosis in the vessel wall,<br>no bleeding or edema | No | mostly lymphocytes, macrophages, pneumocytes, isolated eosinophils, often karyorrhexis | No | No |
| 49 | MOCK | approximately 50% dense areas, clearly bronchial-associated | Bronchitis (lymphocytes and granulocytes in the lumen), bronchial epithelium sporadically with inflammatory infiltration, little damage. | Vascular walls with inflammatory infiltration,<br>multiple hemorrhages in the tissue,<br>2 x edema around vessels | No | mostly lymphocytes, macrophages, pneumocytes,<br>no granulocytes, eosinophils not increased,<br>Foci with massive karyorrhexis | No | No |
| 50 | MOCK | approximately 33% dense areas, clearly bronchial-associated | Bronchitis (lymphocytes and granulozytes in the lumen), bronchial epithelium with little inflammatory infiltration and damage (karyorrhexis). | Vascular walls with inflammatory infiltration (also eosinophils),<br>multiple hemorrhages in the tissue,<br>2 x edema around vessels | No | mostly lymphocytes, macrophages, pneumocytes, isolated eosinophils, foci with karyorrhexis | No | No |
| 51 | MOCK | approximately 33% dense areas, clearly bronchial-associated | Bronchitis (lymphocytes and granulocytes in the lumen), bronchial epithelium sporadically with inflammatory infiltration, little damage. | Vascular walls with inflammatory infiltration (also eosinophils),<br>multiple hemorrhages in the tissue,<br>2 x edema around vessels | No | mostly lymphocytes, macrophages, pneumocytes, isolated eosinophils, frequent karyorrhexis | No | No |
| 52 | MOCK | approximately 33% dense areas (macroscopic), bronchial- and vessel-associated, diffuse distribution | Bronchitis (only few lymphocytes and granulocytes in the lumen) bronchial epithelium with inflammatory infiltration,<br>little damage | Vascular walls with inflammatory infiltration,<br>clear single cell necrosis in the vessel wall,<br>several hemorrhages in the tissue. | No | diffused distributed,<br>mostly lymphocytes, macrophages, pneumocytes,<br>Isolated eosinophils, frequent karyorrhexis | No | No |

| Animal-No. | Vaccine group | % Dense Area | Bronchia | Vessels | DIC | Dense Area | Fibrosis | Syncytia |
|---|---|---|---|---|---|---|---|---|
| 53 | Alum+S | approximately 33% dense areas (macroscopic), bronchial- and vessel-associated, diffuse distribution | Bronchitis (only few lymphocytes, granulocytes and little karyorrhexis in the lumen) bronchial epithelium with little inflammatory infiltration, marked proliferation of the bronchial epithelium, clear damage | Vessel walls only sporadically infiltrated with inflammation, hemorrhages in the tissue | No | mostly lymphocytes, macrophages, pneumocytes, massive eosinophils, frequent karyorrhexis | No | No |
| 54 | Alum+S | 30% dense areas, bronchial- and vessel-associated | Bronchitis (lymphocytes and granulocytes in the lumen), bronchial epithelium: foci of inflammatory infiltration, marked proliferation of the bronchial epithelium, clear damage | Vessel walls only very slightly infiltrated, several hemorrhages in the tissue. | No | mostly lymphocytes, macrophages, pneumocytes, massive eosinophils, frequent karyorrhexis | No | Yes |
| 55 | Alum+S | 40% dense areas, not bronchial- or vessel-associated | Bronchitis (Lymphocytes and granulocytes in the lumen), bronchial epithelium: foci of inflammatory infiltration, marked proliferation of the bronchial epithelium, clear damage | Vascular walls with inflammatory infiltration, significant single cell necrosis in the vascular wall, 1 x hemorrhages in the tissue, 1 x edema around vessels. | No | mostly lymphocytes, macrophages, pneumocytes, massive eosinophils, frequent karyorrhexis | No | Suspicion |
| 56 | Alum+S | 80% dense areas, not clearly bronchial or vessel associated. | Bronchitis (lymphocytes and granulocytes in the lumen), bronchial epithelium: foci of inflammatory infiltration, marked proliferation of the bronchial epithelium, clear damage | Vascular walls with inflammatory infiltration, isolated single cell necrosis in the vascular wall, multiple hemorrhages in tissue, slight edema around vessels. | No | mostly lymphocytes, macrophages, pneumocytes, massive eosinophils, only sporadic karyorrhexis | No | No |
| 57 | Alum+S | 50% dense areas, not clearly bronchial or vessel associated. | Bronchitis (lymphocytes, granulocytes and karyorrhexis in the lumen), bronchial epithelium with very little inflammatory infiltration, no karyorrhexis, little damage | Vascular walls with inflammatory infiltration, isolated single cell necrosis in the vascular wall, sporadic hemorrhages in the tissue. | No | mostly lymphocytes, macrophages, pneumocytes, massive eosinophils, only sporadic karyorrhexis | No | No |
| 58 | Alum+S | 40% dense areas, not clearly bronchial or vessel associated. | Bronchitis (lymphocytes, granulocytes and karyorrhexis in the lumen), bronchial epithelium: Foci of inflammatory infiltration, very few karyorrhexis, little damage | Vascular walls with inflammatory infiltration, 1 x single cell necrosis in the vascular wall, 2 x hemorrhages in the tissue. | No | mostly lymphocytes, macrophages, pneumocytes, massive eosinophils, only sporadic karyorrhexis | No | No |

| Animal-No. | Vaccine group | % Dense Area | Bronchia | Vessels | DIC | Dense Area | Fibrosis | Syncytia |
|---|---|---|---|---|---|---|---|---|
| 59 | MeV | 50% dense areas, not clearly bronchial or vessel associated, diffuse distribution | Bronchitis (lymphocytes, granulocytes and karyorrhexis in the lumen), bronchial epithelium with focal inflammatory infiltration, no karyorrhexis, little damage | Vascular walls with inflammatory infiltration, sporadic single cell necrosis in the vascular wall, multiple hemorrhages in the tissue. | No | mostly lymphocytes, macrophages, pneumocytes, few eosinophils, more neutrophils | No | No |
| 61 | MeV | 40% dense areas, bronchial associated, diffuse distribution | Bronchitis (lymphocytes, granulocytes and karyorrhexis in the lumen), bronchial epithelium: foci of inflammatory infiltration, no karyorrhexis, little damage | Vascular walls with inflammatory infiltration, sporadic single cell necrosis in the vascular walls, edema around the vascular wall, multiple hemorrhages in the tissue. | No | mostly lymphocytes, macrophages, pneumocytes, no eosinophils, more neutrophils sporadic karyorrhexis | No | No |
| 62 | MeV | approximately 20% dense areas (macroscopical), bronchial and vessel associated, diffuse distribution | Bronchitis (lymphocytes, granulocytes and karyorrhexis in the lumen),<br>Bronchial epithelium: foci of inflammatory infiltration,<br>no karyorrhexis, little damage | Vascular walls with inflammatory infiltration,<br>sporadic single cell necrosis in the vascular walls,<br>multiple hemorrhages in the tissue. | No | mostly lymphocytes, macrophages, pneumocytes, eosinophile involved, sporadic karyorrhexis | No | No |
| 63 | MeV | 50% dense areas, not clearly bronchial or vessel associated, diffuse distribution | Bronchitis (lymphocytes and karyorrhexis in the lumen), bronchial epithelium: foci of inflammatory infiltration, sporadic karyorrhexis, proliferation? | Vascular walls with inflammatory infiltration,<br>sporadic single cell necrosis in the vascular walls,<br>edema around the vascular walls, several hemorrhages in the tissue. | No | mostly lymphocytes, macrophages, pneumocytes, eosinophile involved, sporadic karyorrhexis | No | No |
| 64 | MeV | 60% dense areas, not clearly bronchial or vessel associated, diffuse distribution | Bronchitis (little in the lumen), bronchial epithelium with focal inflammatory infiltration, proliferation? | Vascular walls with inflammatory infiltration (especially veins), several hemorrhages in the tissue. | No | mostly lymphocytes, macrophages, pneumocytes, no eosinophils,<br>sporadic karyorrhexis | No | No |
| 65 | MeV-S | 20% dense areas (microscopic), clearly bronchial associated | Bronchitis (little in the lumen), bronchial epithelium hardly affected. | Vascular walls with very little inflammatory infiltration,<br>2 x hemorrhages in the tissue. | No | mostly lymphocytes, macrophages, pneumocytes, isolated eosinophils,<br>no karyorrhexis (1 x ?) | No | No |
| 66 | MeV-S | 10% dense areas, clearly bronchial associated | Bronchitis (little in the lumen), bronchial epithelium hardly affected. | Vascular walls without inflammatory infiltration,<br>several hemorrhages in the tissue. | No | mostly lymphocytes, macrophages, pneumocytes, isolated eosinophils,<br>sporadic karyorrhexis | No | No |
| 67 | MeV-S | 25% dense areas, clearly bronchial associated | Mild bronchitis (mostly little in the lumen), bronchial epithelium with sporadic inflammatory infiltration, otherwise not affected. | Vascular walls with infiltration of eosinophils,<br>several hemorrhages in the tissue. | No | mostly lymphocytes, macrophages, pneumocytes, massive eosinophils around vessels,<br>no karyorrhexis | No | No |
| 68 | MeV-S | 50% dense areas, clearly bronchial-associated | Mild bronchitis (mostly little in the lumen), bronchial epithelium with focal inflammatory infiltration otherwise not affected. | Vascular walls infiltration of eosinophils,<br>several hemorrhages in the tissue. | No | mostly lymphocytes, macrophages, pneumocytes, massive eosinophils around vessels,<br>no karyorrhexis | No | No |

| Animal-No. | Vaccine group | % Dense Area | Bronchia | Vessels | DIC | Dense Area | Fibrosis | Syncytia |
|---|---|---|---|---|---|---|---|---|
| 69 | MeV-S | 30% dense areas, clearly bronchial associated | Mild bronchitis (lymphocytes and karyorrhexis in the lumen), bronchial epithelium with focal inflammatory infiltration, significant damage to the bronchial epithelium | Vascular walls with inflammatory infiltration,<br>karyorrhexis in the vascular wall,<br>edema around vessels,<br>massive hemorrhage in the tissue. | No | mostly lymphocytes, macrophages, pneumocytes, significant amount of eosinophils,<br>foci with karyorrhexis | No | Yes |
| 70 | MeV-S | 60% dense areas, not clearly bronchial associated | Mild bronchitis (lymphocytes, karyorrhexis in the lumen), bronchial epithelium with focal inflammatory infiltration; significant damage to the bronchial epithelium. | Vascular walls with inflammatory infiltration,<br>edema in the vascular wall,<br>edema around the vessels,<br>massive hemorrhages in the tissue. | No | mostly lymphocytes, macrophages, pneumocytes,<br>isolated eosinophils,<br>sporadic karyorrhexis | No | No |

**Supplementary Table 2. Histopathological analysis of lung tissue of vaccinated Syrian hamsters batch #2 upon challenge with SARS-CoV-2.** Hamsters were vaccinates with medium (MOCK), $MV_{vac2}$-ATU(P) as measles-only vector control (MeV), $MeV_{vac2}$-SARS2-S(H) (MeV-S), or Alum-adjuvanted Spike protein (Alum+S). The left lobe of vaccinated hamster lungs was dissected 4 dpi. H& E staining revealed histopathological changes and immune cell infiltration assessed by a trained pathologist in a blinded manner. DIC, disseminated intravascular coagulation**.**

| Animal-No. | Vaccine group | % Dense Area | Bronchia | Vessels | DIC | Dense Area | Fibrosis | Syncitia |
|---|---|---|---|---|---|---|---|---|
| #93 | MOCK | about 30% dense areas, mostly bronchial associated | Bronchial epithelium with inflammatory infiltration, granulocytes in the lumen, partial erythrocytes, some bronchial epithelium in the surrounding tissue | Vascular walls sporadically infiltrated with inflammation, marked hemorrhages | No | few densed areas infiltrated with macrophages, pneumocytes, lymphocytes and granulocytes, no eosinophils, moderate karyorrhexis | No | No |
| #98 | MOCK | max. 10%, small foci, not bronchial associated | slightly altered; low inflammatory infiltration, some inflammatory cells in the lumen | prominent endothelium, minor hemorrhage | No | with increased eosinophils; macrophages and lymphocytes | No | No |
| #105 | MOCK | max. 10%, in a single condensed area, not bronchial associated | slightly altered; low inflammatory infiltration underneath the epithelium | prominent endothelium, minor hemorrhage | No | 1 x dense area with macrophages and lymphocytes, hardly any granulocytes, no eosinophils. moderate karyorrhexis. | No | No |
| #109 | MOCK | about 30%, dense areas not clearly bronchial associated | slightly altered; epithelium sporadically with inflammatory infiltration; granulocytes in the lumen | prominent endothelium, hemorrhage | No | Macrophages, pneumocytes, lymphocytes, hardly any granulocytes and eosinophils. Isolated foci with karyorrhexis. | No | No |
| #113 | MOCK | max. 10%, in a single condensed area, | Epithelium clearly altered; scattered inflammatory infiltration, no granulocytes in the lumen | prominent endothelium, significant bleeding | No | Macrophages, pneumocytes, lymphocytes, hardly any granulocytes and eosinophils. Isolated foci with karyorrhexis. | No | No |
| #91 | MeV-S | max. 10%, clear relation to bronchi. | Marked proliferation of the epithelium into the surrounding tissue. Epithelium with inflammatory infiltration | Prominent endothelium, minor hemorrhage? | No | Some eosinophils around vessels, macrophages, pneumocytes, lymphocytes and granulocytes, hardly any karyorrhexis. | No | No |
| #95 | MeV-S | 10% dense areas, small multiple foci, bronchial associated | Epithelium not signifiantly altered. | Prominent endothelium, scattered karyorrhexis in vessel walls, only 1 x hemorrhage | No | Some eosinophils in dense area, macrophages, pneumocytes, lymphocytes and granulocytes, hardly any karyorrhexis, | No | No |
| #99 | MeV-S | minimal; only 3 small foci, strictly bronchial associated | Epithelium not significantly altered. Scattered Epithelium in the surrounding tissue. | Prominent endothelium, no hemorrhage | No | Some eosinophils in dense area, macrophages, pneumocytes, lymphocytes and granulocytes, no karyorrhexis. | No | No |

| Animal-No. | Vaccine group | % Dense Area | Bronchia | Vessels | DIC | Dense Area | Fibrosis | Syncitia |
|---|---|---|---|---|---|---|---|---|
| #102 | MeV-S | minimal; only smallest foci, bronchial associated | Regular | Prominent endothelium, no hemorrhage | No | Some eosinophils in dense area, macrophages, pneumocytes, lymphocytes and granulocytes, no karyorrhexis. | No | Suspected |
| #114 | MeV-S | 30% dense areas, bronchial associated | Bronchitis, lymphocytes and a few granulocytes in the lumen; inflammatory infiltration of the epithelium. Epithelium in the surrounding tissue. | Prominent endothelium, perivasculitis with some eosinophils, hemorrhage in the surrounding area. | No | Macrophages, pneumocytes, lymphocytes. Hardly any granulocytes and eosinophils. Sporadic karyorrhexis. | No | No |
| #94 | Alum+S / Dexa | 10% dense areas, not bronchial associated | Bronchitis, epithelium with inflammatory infiltration | Vasculitis, prominent endothelium, low inflammatory infiltration of the vessel walls | No | Some eosinophils especially around vessels (poorly visible due to hemorrhages) Macrophages, pneumocytes, lymphocytes and granulocytes. | No | No |
| #97 | Alum+S | 50% dense areas, not bronchial associated | Bronchitis, epithelium with inflammatory infiltration, granulocytes and lymphocytes in the lumen, bronchial epithelium in the surrounding tissue | Marked vasculitis with inflammatory infiltration of the wall (incl. eosinophils), many eosinophils in the surrounding of the vessel walls | No | Many eosinophils especially around vessels. Macrophages, pneumocytes, lymphocytes and granulocytes. | No | No |
| #101 | Alum+S | 50% dense areas, not bronchial associated | Epithelium not infiltrated by inflammation, blood in the lumen | Marked vasculitis with inflammatory infiltration of the vessel wall, no edema around vessels, massive hemorrhage in the surrounding tissue. | No | Massive foci of eosinophils (vessels?), macrophages, pneumocytes, lymphocytes and granulocytes. | No | No |
| #104 | Alum+S | 60% dense areas, not bronchial associated | Epithelium little affected; granulocytes in the lumen | Marked vasculitis with inflammatory infiltration of the wall. | No | Massive foci of eosinophils (vessels?), macrophages, pneumocytes, lymphocytes and granulocytes. | No | No |
| #106 | Alum+S | 70% dense areas, not bronchial associated | Bronchitis, epithelium with little inflammatory infiltration, epithelium irregular, granulocytes (eosinophils) in the lumen | Marked vasculitis with low infiltration of eosinophils in the vessel wall, massive hemorrhage in the surrounding tissue. | No | Many eosinophils especially around vessels. Macrophages, pneumocytes, lymphocytes and granulocytes. Karyorrhexis. | No | No |
| #107 | Alum+S / Dexa | 30% dense areas, not bronchial associated | Bronchitis, epithelium with little inflammatory infiltration, epithelium irregular, granulocytes (eosinophils) in the lumen | Marked vasculitis with inflammatory infiltration of the vessel wall, hemorrhage in the surrounding area. | No | Some eosinophils, but more granulocytes, macrophages, pneumocytes, lymphocytes. | No | No |
| #111 | Alum+S / Dexa | 20% dense areas, bronchial associated | Epithelium hardly affected, single eosinophils in lumen | Prominent endothelium, hemorrhages. | No | Mild Infektion, Eosinophils present, often focal, Macrophages, pneumocytes, lymphocytes, granulocytes, karyorrhexis. | No | No |

| Animal-No. | Vaccine group | % Dense Area | Bronchia | Vessels | DIC | Dense Area | Fibrosis | Syncitia |
|---|---|---|---|---|---|---|---|---|
| #112 | Alum+S | 60% dense areas, not bronchial-associated, highly condensed | Epithelium hardly affected, Detritus and granulocytes in the lumen | Prominent endothelium, sporadic inflammatory inflammation in the vessel wall, massive perivascular lymphatic infiltration. | No | Some eosinophils, granulocytes, macrophages, pneumocytes, lymphocytes low karyorrhexis. | No | No |
| #116 | Alum+S / Dexa | minimal; only smallest foci, strictly bronchial associated | Regular | Prominent endothelium, sporadic inflammatory infiltration. | No | Eosinophils clearly present, macrophages, pneumocytes, lymphocytes. | No | No |
| #117 | Alum+S / Dexa | 20% dense areas, not clearly bronchial associated | Epithelium appears disorganized; no infiltration of epithelium | Prominent endothelium, vessel wall infiltrated with eosinophils. Massive eosinophils in lumen + surrounding tissues. | No | Many eosinophils, macrophages, pneumocytes, lymphocytes, granulocytes. | No | No |
| #118 | Alum+S | 70% dense areas, not bronchial associated, highly condensed | Epithelium appears disorganized; no infiltration of epithelium low amount of inflammatory cell in the lumen | Prominent endothelium massive lymphatic sheath, massive eosinophils in the lumen + around the vessels, hemorrhages. | No | Massive eosinophils present, macrophages, pneumocytes, lymphocytes, granulocytes. | No | No |
| #120 | Alum+S / Dexa | 40% dense areas, not bronchial associated | Epithelium clearly altered; occasional inflammatory infiltration, granulocytes in the lumen; epithelium in the surrounding tissue | Prominent endothelium, inflammatory infiltration (eosinophils) of the wall, no hemorrhage. | No | Some eosinophils, granulocytes, macrophages, pneumocytes, lymphocytes low karyorrhexis. | No | Suspected |
| hamster #1 | naive | Regular | Regular | Regular. | No | Regular. | No | No |
| hamster #2 | naive | Regular | Regular | Regular. | No | Regular. | No | No |

**Supplementary Table 3**

Primer and Probe sets used for quantitative qRT-PCR. 6- Carboxyfluorescein (6FAM), BlackBerry® Quencher (BBQ), Cyanine 5 (Cy5)

| Primer/Probe name | Reference | Sequence |
|---|---|---|
| E_Sarbeco_F | Corman et al. 2020 | 5´-ACAggTACgTTAATAgTTAATAgCgT-3´ |
| E_Sarbeco_R | Corman et al. 2020 | 5´-ATATTgCAgCAgTACgCACACA-3´ |
| E_Sarbeco probe | Corman et al. 2020 | 5‘-(6FAM)-ACACTAgCCATCCTTACTgCgCTTCg(BBQ)-3‘ |
| Il-5_For | Mendlovic et al. 2014 | 5’-gCCgTAgCCATggAgATC-3’ |
| IL-5 Seq Rev | this work | 5’-CgATgCACAgCTggTgCT-3’ |
| IL-5 probe | this work | 5’-(Cy5)-AgCTgTCCACTCACCgAGCTCTACTGAC (BBQ)-3` |
| RPL18 F | Zivec et al. 2011 | 5‘-gTTTATgAgTCgCACTAACCg-3‘ |
| RPL18 R | Zivec et al. 2011 | 5‘-TgTTCTCTCggCCAggAA-3‘ |
| RPL18 probe | Zivec et al. 2011 | 5‘-(Cy5)-TCTgTCCCTgTCCCggATgATC(BBQ)-3‘ |
| Eotaxin forward | Stanelle-Bertram et al. | 5’- AgAgAgCCTgAgACCAACAC-3’ |
| Eotaxin reverse | Stanelle-Bertram et al. | 5’-AACTgggATAgAgCCTgggTg-3’ |
| Eotaxin-probe | this work | 5`-(6FAM)-TTgTggCCACTgCCTTCACCTC (BBQ)-3’ |
| IL-4-F | Espitia et al. 2010 | 5'-ACAgAAAAAgggACACCATgCA-3' |
| IL-4-R | Espitia et al. 2010 | 5'-gAAgCCCTgCAgATgAggTCT-3' |
| IL-4 probe | Espitia et al. 2010 | 5'-(6FAM)-AgACgCCCTTTCAgCAAggAAgAACTCC-(BBQ)-3' |
| IL-13-F | Espitia et al. 2010 | 5'-AAATggCgggTTCTgTgC-3' |
| IL-13-R | Espitia et al. 2010 | 5'-AATATCCTCTgggTCTTgTAgATgg-3' |
| IL-13 probe | Espitia et al. 2010 | 5'-(Cy5)-TggATTCCCTgACCAACATCTCTAgTTgC (BBQ)-3' |
